# Interpretable variant effect prediction from genomic foundation model representations

**DOI:** 10.64898/2026.04.10.717844

**Authors:** Michael T. Pearce, Thomas Dooms, Ryo Yamamoto, Shant Ayanian, Alexander J. Ryu, Joshua Meehl, Carl Molnar, Tuomo Kiiskinen, Mark Bissell, Dron Hazra, Ching Fang, Nam Nguyen, Michael Anderson, Collin Osborne, Patrick Duffy, Bridget Toomey, Elena Myasoedova, Eric Klee, Panos Korfiatis, Matt Redlon, Archa Jain, Daniel Balsam, Nicholas K. Wang

## Abstract

Scientific foundation models learn high-dimensional representations from diverse data modalities, yet what they encode and how to extract that knowledge remain open questions. Here we show that probing the internal representations of Evo 2, a 7-billion-parameter genomic foundation model, enables accurate and interpretable genetic variant effect prediction. We introduce a covariance-based probe that captures second-order structure from Evo 2 sequence embeddings to predict variant pathogenicity across variant types and functional consequences, matching or exceeding specialized predictors within their domains. To ground these predictions in known biological mechanisms, we train a complementary panel of probes on existing annotations to detect which genomic properties are disrupted by a variant. This categorized evidence is then integrated with each variant’s genomic context through a language model to generate variant-specific mechanistic hypotheses. Our pathogenicity predictions correlate with experimental measures of variant function, clinical penetrance, and biobank disease associations while the mechanistic hypotheses are consistent with expert reviews, known mechanism classes, and downstream molecular readouts. We release pathogenicity scores, disruption profiles, and contextualized interpretations for 4.2 million variants from the ClinVar database as an open resource through the Evo Variant Effect Explorer (EVEE). More broadly, this structured probing approach offers a general framework for interrogating foundation models across scientific disciplines and grounding their outputs in existing domain concepts.

## Main

Scientific foundation models trained on biological sequence data encode substantial information about the systems they represent [1, 2], yet two questions remain central to their translational utility: what have these models learned, and how can that knowledge be extracted in a form that is both predictive and interpretable? Few problems test this question more directly, or carry higher stakes, than genomic variant effect prediction. The task requires estimating whether a genetic variant causes disease and identifying its biological mechanism to support clinical interpretation. This remains a major bottleneck in genomic medicine, as DNA sequencing routinely surfaces more variants than can be interpreted [3, 4] with most ambiguously classified as Variants of Uncertain Significance (VUS) [5, 6].

Despite a proliferation of variant-effect predictors—protein language models (EVE [7], ESM2 [1], AlphaMissense [8]), supervised sequence-to-function models (Enformer [9], Borzoi [10], Nucleotide Transformer v3 [11, 12], AlphaGenome [13]), evolutionary DNA models (GPN-MSA [14], Evo 2 [2]), and classical meta-predictors (CADD [15])—two related limitations persist. First, they are fragmented across biological regimes: protein-based methods are largely restricted to missense substitutions, sequence-to-function models focus primarily on regulatory effects, and genome-wide meta-predictors compress heterogeneous evidence into a single score [13, 15, 16]. Second, most methods provide a prediction without a mechanism. For clinical use, a score is rarely sufficient on its own; variant classification relies on categorized evidence for why a variant is damaging, as formalized in frameworks such as the ACMG/AMP guidelines [6].

These limitations need not be intrinsic to foundation models. A single genome-wide model represents coding and non-coding sequence alike in one shared space suggesting the potential to unify prediction across variant classes. The challenge lies in how to query these signals. Variant effects are commonly inferred from output likelihoods: the change in probability a model assigns to a sequence when a variant is introduced. This is powerful, but it collapses the internal state of multi-billion-parameter networks into a single score with documented failure modes [2, 17]. The internal representations are far richer. They capture evolutionary constraints across the genome and organize sequences into interpretable biological features, including gene structure and functional protein elements [2, 12, 18]. We therefore asked whether systematically probing these representations could recover predictive power beyond likelihood scoring and the mechanistic information absent from scalar predictors.

Here we show that structured probing of Evo 2, a 7-billion-parameter genomic foundation model, answers this question: its representations yield variant-effect predictions that are simultaneously accurate and biologically interpretable. A covariance probe reads second-order structure from the embeddings to predict pathogenicity across variant classes, while a complementary panel of annotation probes decodes which biological features a variant disrupts. We leverage large language models to integrate these specific signals with each variant’s unique genomic context to generate testable mechanistic hypotheses. Accuracy and interpretability thus emerge as complementary readouts of the same learned representations rather than competing objectives.

## Results

### A single covariance probe matches or exceeds specialized predictors across variant types

To predict pathogenicity, we probed Evo 2 representations directly, passing each variant’s reference and alternate sequences through the model and computing the difference in per-position embeddings. Rather than mean-pooling these differences, we compute a learned low-rank covariance of the embedding differences from both sequence directions, capturing second-order structure—feature co-occurrences across positions—that mean pooling discards [19–21] (Figure 1a; Supplementary Figures S1–S3). The resulting predictions are well calibrated across 833,970 singlenucleotide variants (SNVs) from ClinVar, a public archive of clinically reported variant classifications (Methods; Supplementary Figure S4). We refer to this calibrated pathogenicity prediction as the EVEE score, after the Evo Variant Effect Explorer (EVEE), the open resource through which we release the system.

**Figure 1.**
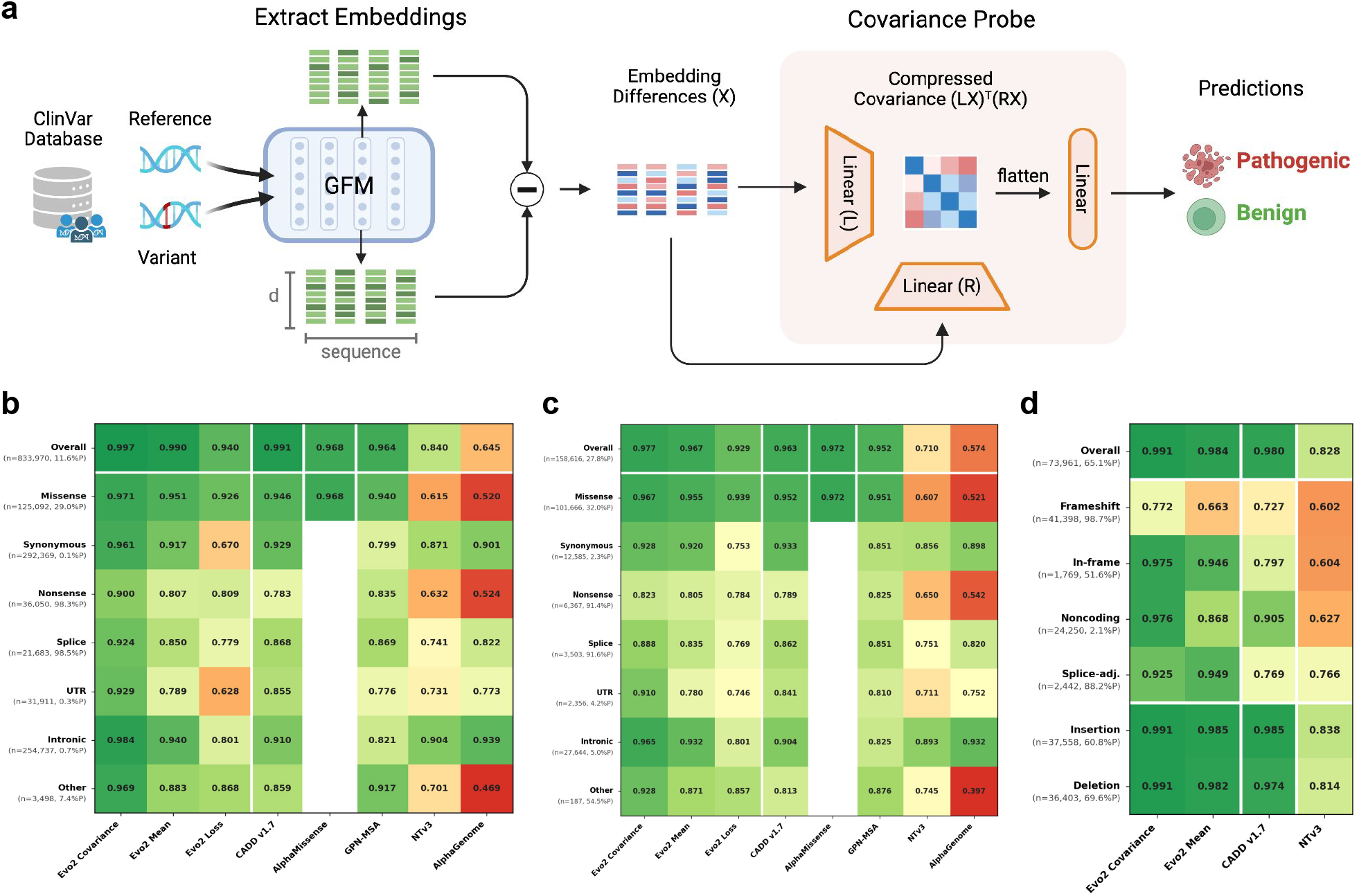
A covariance probe on Evo 2 embeddings predicts pathogenicity across variant consequence types and generalizes zero-shot to indels. (**a**) Experimental design: Evo 2 produces per-position embeddings of reference and alternate sequences; a compressed covariance probe on the embedding difference predicts pathogenicity. (**b**) ClinVar SNV pathogenicity AUROC across consequence types (833,970 variants, ≥1-star review). Blank cells: methods without predictions for that variant type. (**c**) AUROC on a deconfounded benchmark (158,616 variants) that balances pathogenic and benign labels within each consequence type, controlling for consequence-type priors. (**d**) Zero-shot AUROC on 73,961 ClinVar indels stratified by consequence type and insertion vs. deletion.

Variant consequence type is a strong determinant of expected pathogenicity rates: in this dataset, 98.3% of nonsense variants are pathogenic, compared with 0.1% of synonymous variants. An aggregate AUROC (0.997 across all SNVs) is therefore largely confounded by a model’s ability to recover consequence-type priors rather than variant-level discrimination [18, 22]. To account for this, we evaluated performance within each consequence type, using gene-level holdout so that no gene was shared between train and test sets (Methods). Stratified this way, our covariance probe discriminates pathogenic from benign variants (ranging from AUROC 0.900 for nonsense variants to 0.984 for intronic; *n* ranges 3,498–292,369; Figure 1b), outperforming Evo 2 loss-based likelihood scoring, CADD, GPN-MSA, NTv3, and AlphaGenome across consequence types and matching AlphaMissense, the missense-specialized comparator, on missense variants. It also outperformed a mean-pooled probe trained on the same Evo 2 embeddings, isolating covariance structure as a major source of the gain.

To further remove consequence-type priors, we evaluated performance on a deconfounded benchmark that balances pathogenic and benign labels within each consequence type (consequence-only accuracy 96% *→* 73%; Supplementary Figure S5). The Evo 2 covariance probe remained competitive, matching or exceeding the strongest available comparator across consequence types (Figure 1c). Finally, although trained only on SNVs, the probe generalized zeroshot to 73,961 indels, a variant class that many existing methods fail to accommodate, outperforming CADD v1.7 across indel types (overall AUROC 0.991 vs. 0.980; Figure 1d). Performance was notably lower for frameshifts (0.772), a weakness likely exacerbated by severe class imbalance (98.7% pathogenic).

### Predictions generalize beyond conservation to held-out functional assays

High accuracy on ClinVar could reflect a coarser signal— evolutionary conservation, which flags positions where variation is rarely tolerated but says little about the specific variant, and predictors that rely heavily on it can lose variant-level resolution where conservation is highest. Across the conservation spectrum (phyloP), the covariance probe maintained high AUROC from fast-evolving to highly conserved sites, whereas CADD and GPN-MSA degraded at the extremes (Figure 2a), indicating that Evo 2 representations encode functional information beyond sequence conservation alone. UMAP visualization of reference-subtracted Evo 2 embeddings further showed that the learned representation space organizes variants by both pathogenicity and consequence type (Figure 2b,c).

**Figure 2.**
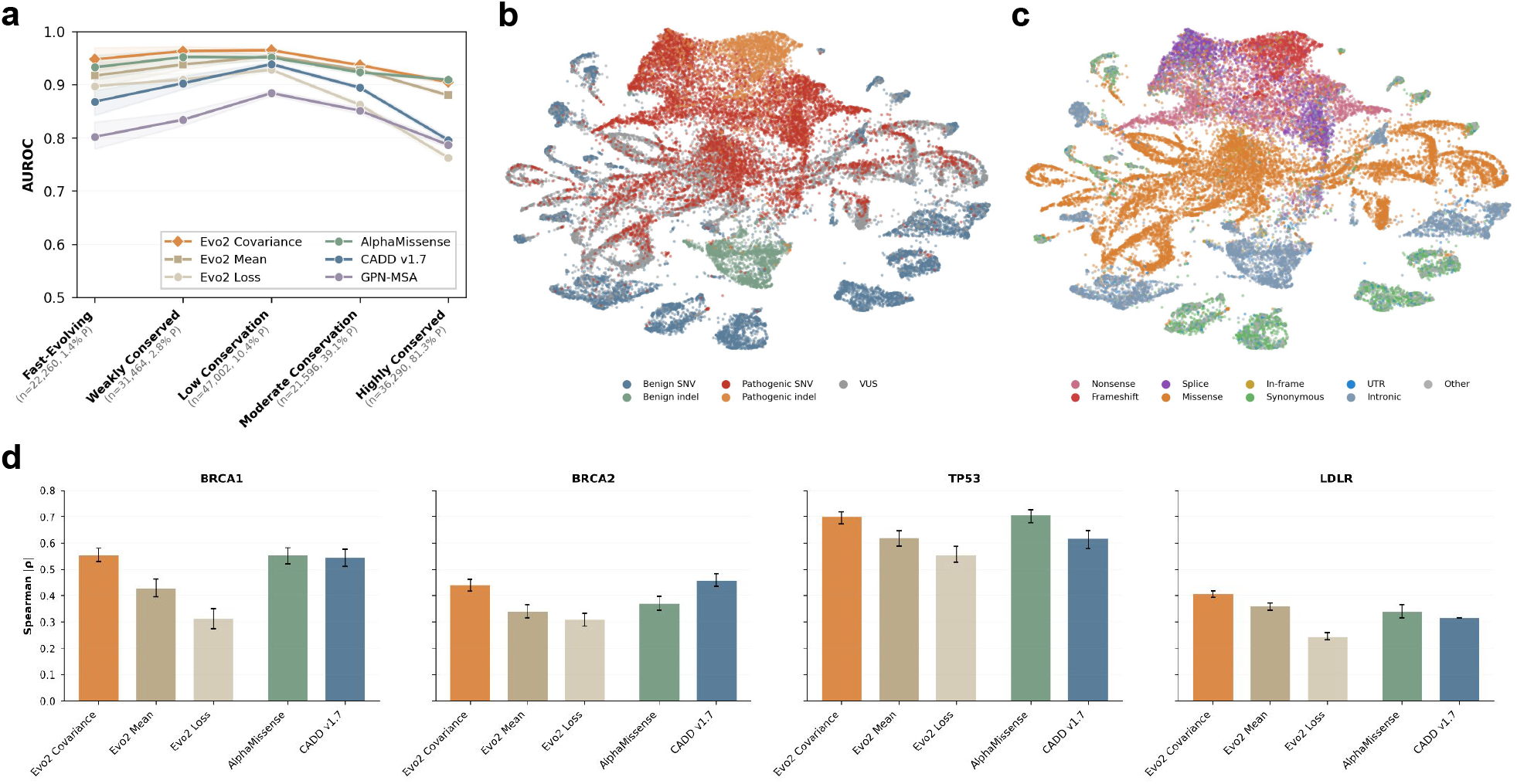
Probe performance holds across the conservation spectrum and transfers to deep mutational scans. (**a**) AUROC by evolutionaryconservation tier (phyloP100way). (**b**,**c**) UMAP of variant covariance embeddings colored by (b) ClinVar pathogenicity label (benign, VUS, pathogenic) and (c) consequence type, for SNVs and indels. (**d**) DMS generalization per gene; Spearman |*ρ*| between predicted scores and continuous DMS readouts. Error bars: 95% bootstrap CI.

As a direct experimental validation beyond ClinVar, we evaluated transfer to deep mutational scan (DMS) datasets— high-throughput assays that measure the functional effect of thousands of variants in a gene—for four genes held out from probe training: BRCA1 [23], BRCA2 [24], TP53 [25], and LDLR [26] (Figure 2d). We measured correlation between model predictions and continuous DMS functional scores (Spearman |*ρ*|), comparing Evo 2 mean-pooled and covariance-pooled probes with CADD, AlphaMissense, and Evo 2 loss-based scoring. Across all four genes, Evo 2 embedding probes outperformed loss-based scoring. The covariance probe was competitive with AlphaMissense and CADD: it led on LDLR (|*ρ*| = 0.41 versus 0.34 for AlphaMis-sense and 0.32 for CADD), matched AlphaMissense on BRCA1 (|*ρ*| = 0.55) and TP53 (| *ρ*| = 0.70), and was comparable to CADD on BRCA2 (|*ρ*| = 0.44 versus 0.46). Covariance probes consistently outperformed mean-pooled probes, corroborating the ClinVar results and confirming that second-order embedding structure captures functional signal lost under mean pooling. Together, these results show that Evo 2 representations generalize from clinical labels to experimental measures of variant effects.

### Disruption profiles recover disease mechanisms beyond pathogenicity

Accurate prediction is most useful when paired with an explanation of what biological properties a variant disrupts. We therefore trained token-level probes on labeled sequences from genomic annotation databases to map known sequence properties onto Evo 2’s latent space. Applied to reference and alternate sequences, these probes measure how a variant changes each property, yielding a disruption profile which we pass to a language model to synthesize into natural-language explanations (Figure 3a). The displayed annotation panel comprises 236 binary probes spanning genic regions, structural domains, protein families, and splice modules, with a median AUROC of 0.919 (Figure 3b).

**Figure 3.**
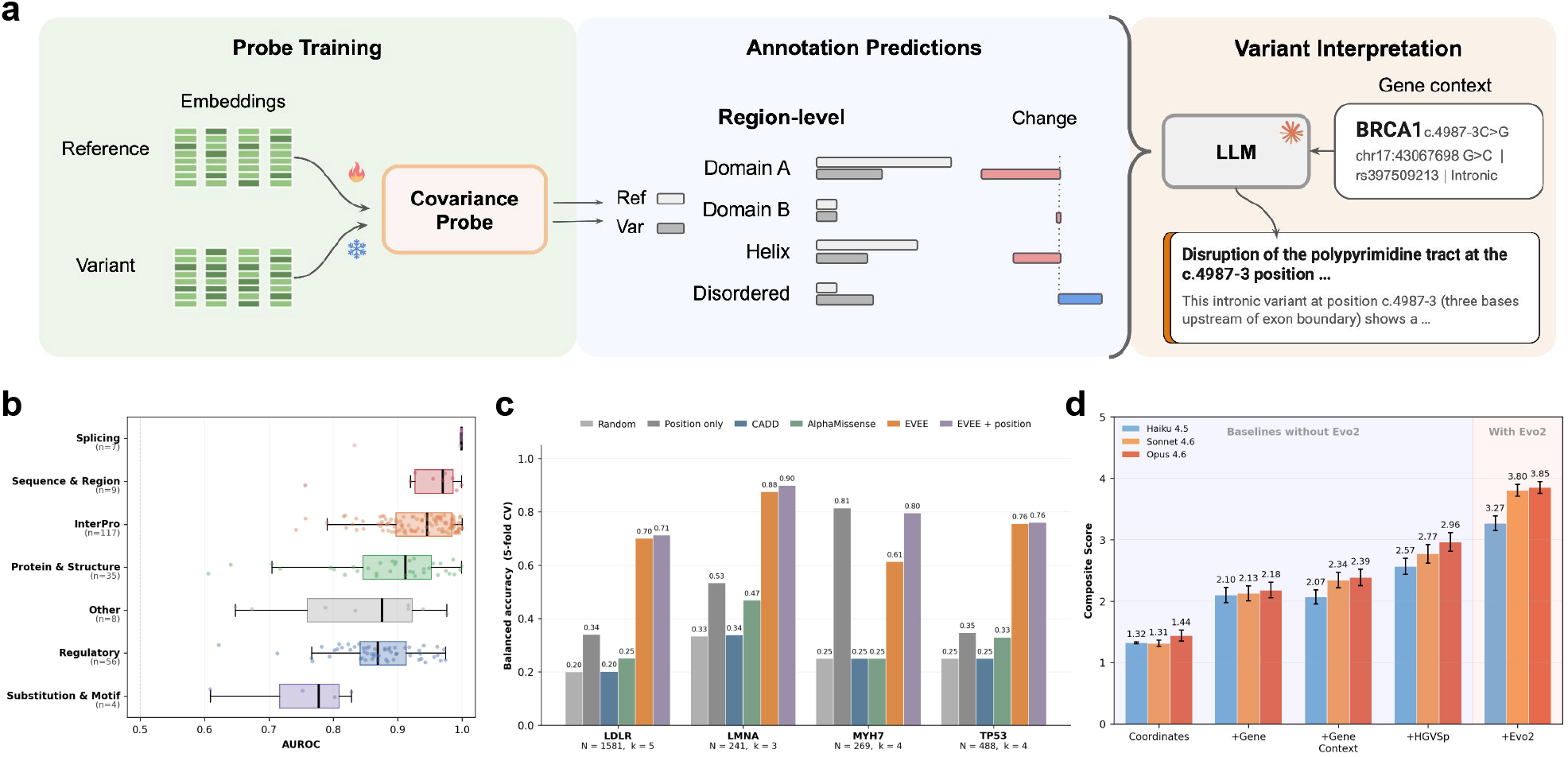
Disruption profiles recover disease-mechanism classes and improve variant interpretation. (**a**) Interpretability framework: annotationprobes trained on Evo 2 reference-sequence embeddings produce per-variant disruptions (Δ = variant − reference) that an LLM synthesizes into naturallanguage explanations. (**b**) Annotation probe AUROC by category (236 binary probes; per-head jitter). (**c**) Within-gene supervised mechanism-class recovery across four genes (5-fold CV on EVEE disruption features). Per-gene class counts: LDLR (*k* = 5, *N* = 1,581); LMNA (*k* = 3, *N* = 241); MYH7 (*k* = 4, *N* =269); TP53 (*k* =4, *N* =488). EVEE alone leads on every gene except MYH7, whose mechanism classes coincide with primary structure. (**d**) Composite interpretation-quality scores across cumulative context configurations for three Claude model tiers (Haiku 4.5, Sonnet 4.6, Opus 4.6); error bars: 95% CI.

These disruption profiles contain mechanistic information beyond position or pathogenicity alone. In allelic series— sets of variants in the same gene that act through distinct disease mechanisms—a logistic-regression classifier using EVEE disruption features discriminated mechanism classes across LDLR, LMNA, MYH7, and TP53 (Figure 3c). The disruption profiles outperformed a position-only baseline (mean balanced accuracy 0.74 versus 0.51), whereas pathogenicity scores alone did not distinguish mechanism classes above chance (mean balanced accuracy 0.33 for AlphaMissense and 0.26 for CADD). Thus, the same foundation-model representations that support pathogenicity prediction also encode separable, biological mechanisms of effect.

To test whether natural-language synthesis of these disruption profiles yields clinically concordant explanations, we scored interpretations for 147 expert-reviewed ClinVar variants (*≥* 3 stars; 98 benign or likely benign, 49 pathogenic or likely pathogenic) on mechanism coverage, biological accuracy, and specificity against expert submission text with an LLM-as-judge. As we progressively added context to the interpreting model, the composite score rose from 1.31 (coordinates only) to 3.80/5 (full context) with Claude Sonnet 4.6, with the largest gain coming from the Evo 2 disruption profiles (+1.03; Figure 3d). Because well-characterized variants may be memorized by the interpreting model and the judge scores only against provided ClinVar evidence, we interpret these scores as a relative measure of the disruption profile’s contribution rather than an absolute benchmark of clinical interpretation quality (Methods; Supplementary Figures S6,S7,S8).

Concretely, we feed the top 10 disruptions and variant metadata (gene, consequence type, and HGVS nomenclature) into an LLM, which generates a contextualized explanation for each variant’s predicted pathogenicity (Figure 4a). For BRCA1 c.5278 *−* 1G*>*A, a variant that destroys the essential splice-acceptor site at the intron 19/exon 20 boundary, the disruption profile shows loss of polypyrimidine-tract (Δ = *−* 0.47) and exon *→* intron-boundary (Δ = *−* 0.44) signal at the canonical acceptor, together with gain of intron-region (Δ = +0.83) and splice-acceptor (Δ = +0.54) signal eight bases into exon 20. This predicts use of a cryptic acceptor site 8 bases downstream. The same framework produces specific mechanisms for nonsense variants such as LDLR p.Ser306Ter, which truncates the protein before the EGF precursor homology domain, and missense variants such as RET p.Gly798Ser, which is predicted to disrupt kinase-fold backbone flexibility (Supplementary Figures S9,S10).

**Figure 4.**
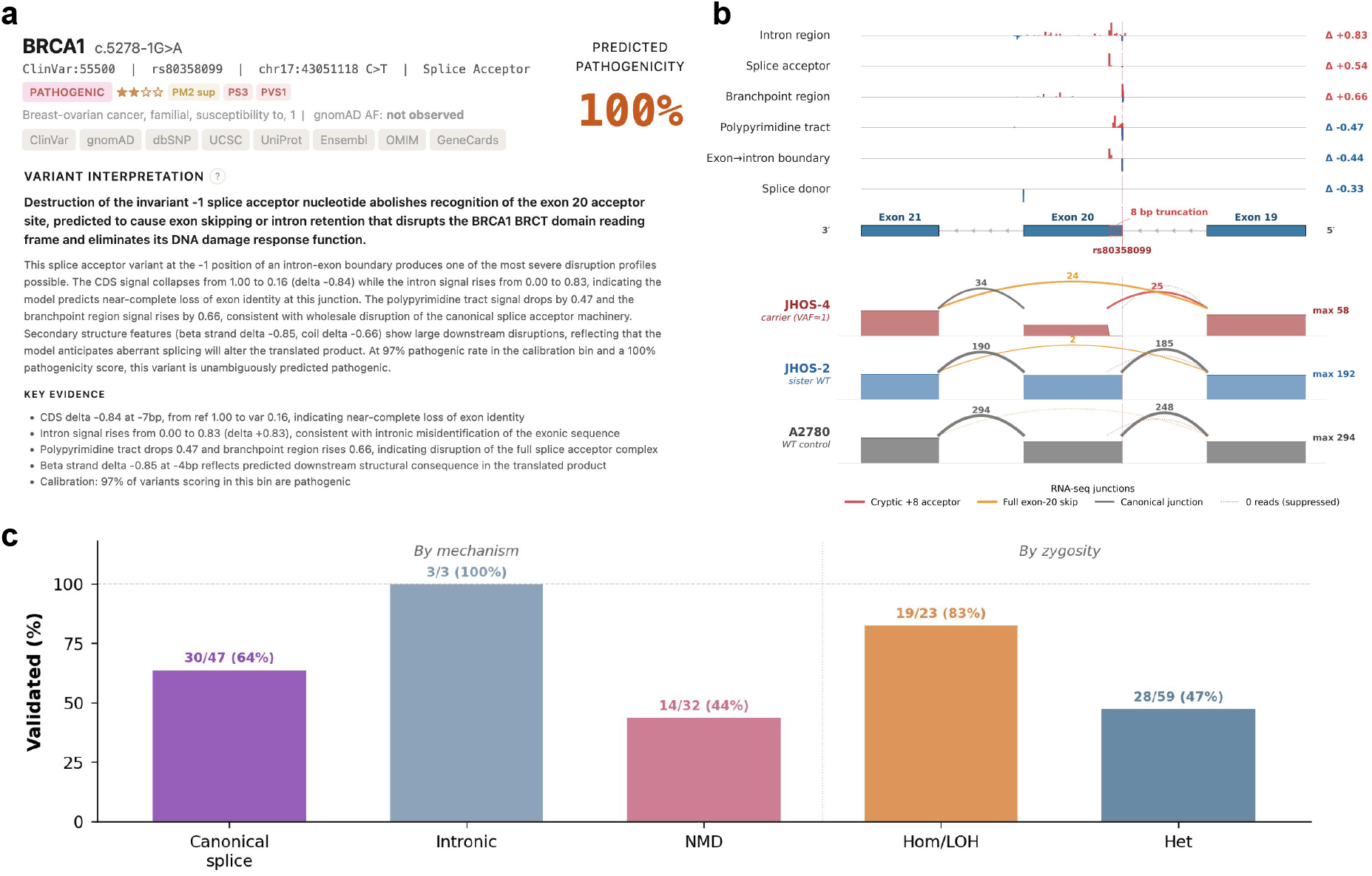
Disruption profiles nominate testable mechanistic hypotheses supported by RNA-seq. (**a**) Example variant interpretation for BRCA1 c.5278-1G*>*A: disruption profile and LLM-synthesized mechanism statement with ACMG-aligned evidence summary. (**b**) RNA-seq validation of the panel-(a) prediction in three CCLE ovarian cancer cell lines (sashimi plots, exons 19–21). JHOS-4 (variant carrier) shows complete loss of canonical exon-20 inclusion,with reads split between the cryptic +8 acceptor (red arcs) and full exon-20 skip (gold arcs); BRCA1 wild-type controls JHOS-2 and A2780 splice canonically (blue arcs). (**c**) Aggregate RNA-seq validation of EVEE-predicted splice, branchpoint, and NMD variants across 82 variant–carrier pairs in CCLE: validation rate stratified by mechanism class and by zygosity (Hom/LOH 19*/*23, 83%; heterozygous 28*/*59, 47%).

We directly validated the BRCA1 c.5278 *−* 1G*>*A prediction using RNA sequencing in three ovarian cancer cell lines from the Cancer Cell Line Encyclopedia (CCLE; Figure 4b). JHOS-4, which carries the variant on essentially all gene copies (variant allele fraction 0.96 by whole-exome sequencing), showed complete loss of canonical exon-20 inclusion, with mutant reads split between full exon-20 skipping and the exact +8 cryptic acceptor that the disruption profile had flagged—matching EVEE’s mechanistic prediction at nucleotide resolution. Two BRCA1 wild-type controls, JHOS-2 and A2780, showed canonical splicing (per-junction read counts and the 8-bp truncated exon-20 region absent from JHOS-4 coverage are reported in Methods).

Beyond this single variant, we scored predicted pathogenic splice, branchpoint, frameshift, and premature-termination-codon (PTC) variants against CCLE RNA-seq across matched carrier cell lines. Of 82 unique variant–carrier pairs assayed, 47 (57%) met the disruption threshold (*>* 70% of mutant-allele reads showing aberrant splicing or nonsense-mediated decay). Validation was higher among carriers in which the variant was present on all gene copies (19*/*23, 83%) than among heterozygous carriers (28*/*59, 47%), as expected due to the wild-type allele diluting mutant RNA-seq signal (Figure 4c). These results show that disruption profiles can generate testable, nucleotide-resolution hypotheses, a subset of which are concordant with independent transcriptomic observations.

### Predictions track clinical penetrance and population-scale disease risk

A pathogenicity score is only useful if it tracks real outcomes—whether a carrier develops disease, and whether variants associate with disease at scale. In Mayo Clinic’s Tapestry cohort, we evaluated 147 unique LDLR variant carriers with expert-curated Familial Hypercholesterolemia (FH; inherited high-LDL-cholesterol disorder) severity tiers. Restricting to carriers on the FH-severity axis—clinical FH (definite or probable; *n* = 111), suspected FH (possible; *n* = 20), and presymptomatic carrier (variant-positive without FH manifestation; *n* = 16)—EVEE scores discriminated FH-manifesting carriers from presymptomatic carriers at AUROC = 0.91 (95% CI 0.84–0.96), compared with 0.70 (0.58–0.82) for AlphaMissense and 0.65 (0.51– 0.79) for CADD (Figure 5a). Median EVEE scores decreased monotonically across severity tiers (clinical FH 0.76, suspected FH 0.10, presymptomatic 0.01; Figure 5b), indicating that the score tracks clinical penetrance rather than ClinVar label alone.

**Figure 5.**
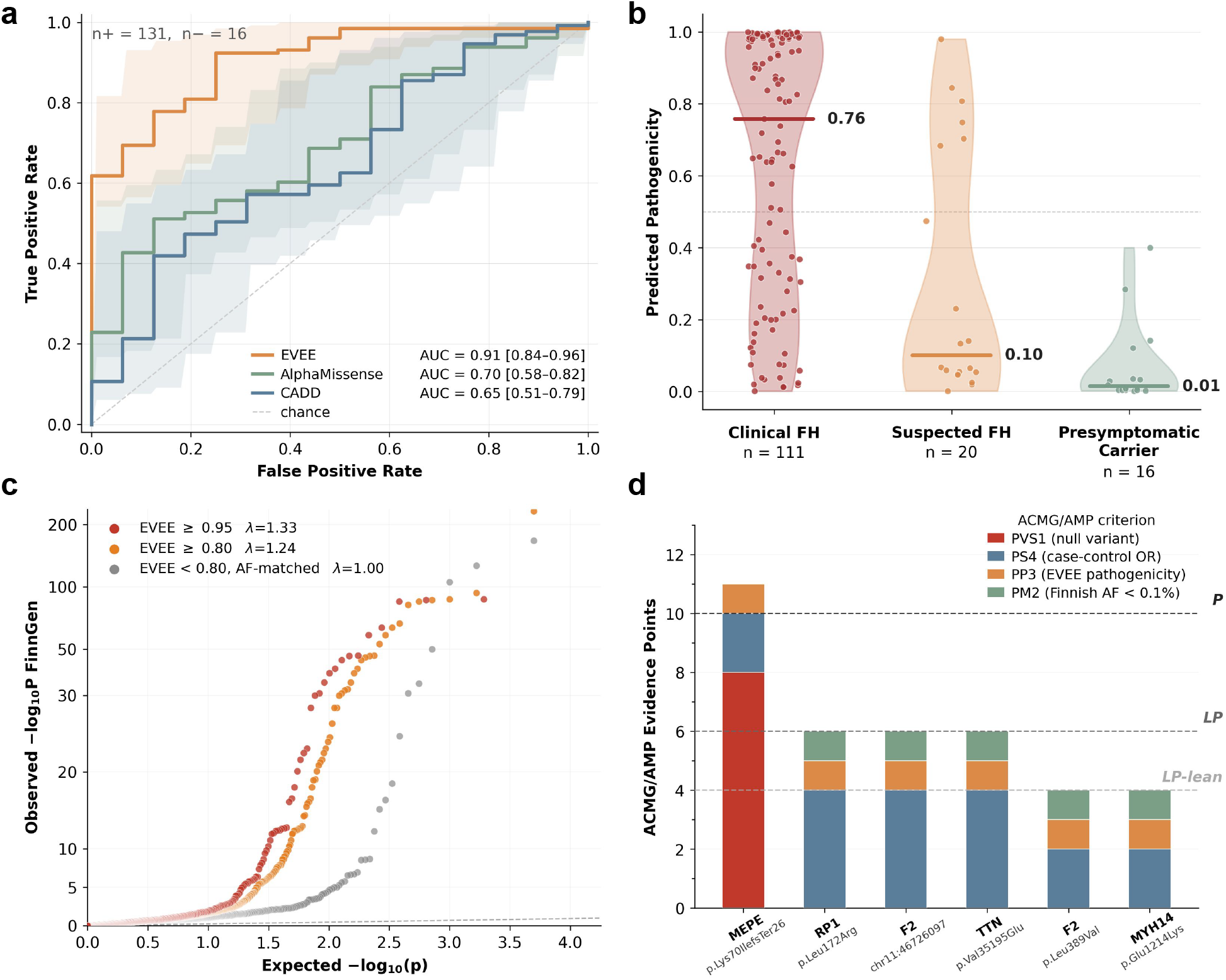
EVEE scores track clinical penetrance, enrich for biobank disease associations, and support ACMG/AMP variant nomination. (**a**) Clinicalpenetrance validation in the Mayo Clinic Tapestry cohort: ROC curve distinguishing FH-manifesting LDLR variant carriers (*n*+ = 131) from presymptomatic carriers (*n*− = 16). Shaded bands and bracketed AUROC values denote 95% CIs (2,000-resample stratified bootstrap). (**b**) Per-tier distribution of the EVEE score across the three FH severity tiers (Clinical FH, Suspected FH, Presymptomatic) in the joined Tapestry × ClinVar LDLR cohort; per-tier medians fall monotonically (0.76 → 0.10 → 0.01). (**c**) Independent validation in FinnGen R12: grouping variants by EVEE score reveals dose-responsive diseaseassociation enrichment in a biobank the model was not trained on. Per-variant QQ plot of FinnGen R12 association *p*-values (minimum over 2,489 endpoints, Bonferroni-corrected) for EVEE score ≥0.95 (*λ*GC =1.33, *n*=958) and the more inclusive ≥0.80 set (*λ*GC =1.24, *n*=2,497), versus an allele-frequency– matched control of variants with EVEE *<* 0.80 (*λ*GC = 1.00; 5:1 AF-matched to the ≥ 0.80 set). *y*-axis is symlog (linear below −log10(*p*) = 30); dashed line: expected null. (**d**) ACMG/AMP structured evidence for the six FinnGen R12 candidates accumulating sufficient evidence under the Tavtigian (2018) point-weighted framework [28] to support ClinVar submission. Stacked bars: per-variant evidence points by criterion (PVS1, PS4, PP3, PM2). Dashed lines: Tavtigian (2018) thresholds (LP-lean ≥ 4, LP ≥ 6, P ≥ 10).

As an early-deployment demonstration, we applied EVEE to a clinical exome cohort of 588 rheumatoid arthritis patients at Mayo Clinic. Among 225 rare variants (allele frequency *≤* 1% in gnomAD) lacking an established ClinVar classification, 14 (6.2%) reached an EVEE score *≥* 0.90, whereas no benign or likely benign variant exceeded that threshold. High-scoring variants concentrated in immuneregulation and folate/methotrexate-pathway genes relevant to the disease (Supplementary Figure S11).

At population scale, we asked whether EVEE predicts disease association in an independent biobank not used for training. We computed per-variant quantile–quantile plots of variant–phenotype association *p*-values from FinnGen R12 [27], a Finnish national biobank with approximately 500,000 participants and 2,489 disease endpoints, grouping variants by EVEE score. Enrichment of strong associations over the null, summarized by the genomic inflation factor *λ*_GC_, increased with EVEE score: variants with a predicted pathogenicity *≥* 0.95 reached *λ*_GC_ = 1.33, and the broader *≥* 0.80 set reached *λ*_GC_ = 1.24. By contrast, an allele-frequency-matched control set of lower-scoring variants (EVEE *<* 0.80) remained at the null (*λ*_GC_ = 1.00; 5:1 matching across 30 log-spaced minor-allele-frequency bins). Because the matched control shares the same genotyping, ancestry, and multiple-testing pipeline, the elevated *λ*_GC_ in high-EVEE groups reflects disease-association signal rather than allele-frequency or population-stratification artifacts. The signal persisted when the EVEE *≥* 0.95 group was restricted to variants without an established ClinVar classification (*λ*_GC_ = 1.17, *n* = 495), indicating that EVEE captures disease-relevant information beyond previously classified variants.

Finally, we tested whether EVEE outputs can be translated into clinically structured evidence in a biobank-discovery setting. We scored FinnGen R12 rare-variant candidates under the ACMG/AMP framework, which classifies variants by tallying weighted evidence codes such as PVS1 for predicted loss of function, PS4 for case enrichment, PP3 for computational support, and PM2 for population depletion. Rare variants (minor allele frequency *<* 0.5% in the Finnish population) showing genome-wide-significant disease association and EVEE score above 0.8 were considered actionable after applying phenotype–gene plausibility and ClinGen gene–disease validity filters (Methods). Of 21 actionable candidates passing initial filters, six accumulated enough evidence under the Tavtigian (2018) point-weighted ACMG/AMP framework [28] to support ClinVar submission (Figure 5d). Beyond ranking variants, EVEE contributes structured evidence that can nominate new candidate disease variants from biobank-scale data.

## Discussion

This work establishes that representations from a genomic foundation model can support both accurate variant effect prediction and mechanistic interpretation. The central result is not simply that a probe improves performance over likelihood scoring, but that both prediction and interpretation can be obtained from the same underlying representations. Our covariance probe reads second-order representation changes to produce accurate pathogenicity estimates across SNVs, indels, coding, splicing, and proximal non-coding effects. The annotation probes read named biological properties from the same representation space, producing disruption profiles that explain how a variant is predicted to act. Together, these results reframe interpretability not as a post hoc compromise on accuracy, but as a way to expose the biological structure that makes the prediction possible.

The approach also defines a general translation layer between specialized scientific foundation models and general-purpose reasoning systems. Evo 2’s embeddings are not themselves interpretable to a clinician or researcher; nor can a language model natively reason over billions of hidden activations. Annotation probes bridge this gap by converting latent representation changes into a compact panel of named, domain-grounded evidence. A language model can then integrate that evidence with variant metadata to generate mechanistic hypotheses aligned with clinical interpretation frameworks. In this sense, the disruption profile functions as an evidence interface, making the foundation model’s internal knowledge legible and usable without requiring the reasoner to access the raw model state.

The validation suite supports the claim that our probes capture true biology rather than ClinVar memorization. Beyond ClinVar benchmarks, EVEE scores correlate with direct experimental measurements in held-out DMS genes, predict penetrance among Mayo Clinic Tapestry LDLR carriers, enrich for disease associations across 2,489 FinnGen endpoints, and nominate six candidate disease variants with sufficient structured ACMG/AMP evidence to support ClinVar submission. The mechanistic layer is likewise experimentally grounded: for BRCA1 c.5278 *−* 1G*>*A, EVEE predicted activation of a cryptic splice acceptor eight bases into exon 20, and RNA-seq in a carrier cell line confirmed the altered splice junction at nucleotide-level resolution. These orthogonal validations are critical because they test whether representation-derived predictions transfer from labels to measurable biological and clinical outcomes.

For all 4.2 million clinically reported ClinVar variants, EVEE provides a pathogenicity prediction, disruption profile, and contextualized interpretation, accessible through a web application (https://evee.goodfire.ai/), bulk download, or a programmatic interface. This resource is intended to support variant prioritization, mechanistic hypothesis generation, and research into how genomic foundation models encode biological information. It is not a substitute for expert review or clinical classification, but it provides structured, inspectable evidence that can be integrated into established interpretation workflows.

Several limitations warrant discussion. Evo 2’s broad evolutionary prior is well suited to many Mendelian disease mechanisms, but may be less complete for regulatory, epistatic, or polygenic effects that lack clear evolutionary signatures. Because the pathogenicity probe is supervised on ClinVar labels, within-ClinVar benchmarks carry a home-domain advantage over predictors not trained on the same label distribution; the DMS, penetrance, RNA-seq, and biobank analyses are therefore the stronger evidence for generalization. The clinical and population-level validations are subject to cohort and ascertainment constraints, including the small presymptomatic-carrier class in Tapestry (*n* = 16) and the use of a single biobank. The annotation probes are limited to known, labeled features; unsupervised methods such as sparse autoencoders applied to genomic foundation-model representations [2, 29, 30] may reveal additional features and could be integrated with disruption profiling. The EVEE-based PP3 criterion is applied at Supporting strength as a heuristic, and formal calibration of evidence strength against observed pathogenicity odds [31] remains necessary. Finally, LLM-generated interpretations should be treated as hypotheses requiring expert review or experimental validation rather than definitive clinical evidence.

As genomic foundation models improve and incorporate additional data types, this framework is designed to accommodate further sources of variation and disease. More broadly, the method is not limited to genomics. Whenever a foundation model is used as a predictor, its internal representations may contain structure that output scores do not expose. Probes make that structure legible, and general-purpose reasoners can integrate the resulting evidence into hypotheses, explanations, and decisions. In genomics, this provides a path from opaque variant scores toward mechanism-aware evidence for clinical interpretation. Across the sciences, it suggests a general strategy for turning foundation-model representations into actionable knowledge.

## Methods

### Genomic foundation model

#### Embeddings

We used the 7B-parameter Evo 2 model [2] to compute (i) per-position log-likelihood scores for variant effect scoring and (ii) intermediate-layer embeddings for representation analysis. Embeddings were extracted from layer 27, selected based on a layer sweep across all 32 blocks (Supplementary Figure S1).

#### Harvesting

We extracted a fixed-size 65k bp window around each mutation site, comprising 49k bp of upstream context and the 16k bp downstream effect. Accuracy increased with window size, indicating a long tail of low-range effects captured by Evo 2 (Supplementary Figure S2). To bound storage, we retained per strand only the embeddings of the 256 positions most divergent between the reference and variant sequences (by cosine similarity). For covariance probes, this top-256 selection performed comparably to keeping a trailing contiguous window after the mutation site (Supplementary Figure S3).

#### Dataset

Both the reference and variant sequences were harvested in the sense and anti-sense directions.For indels, reference and variant sequences were aligned and unmatched positions discarded. Extraction used ~ 20,000 H100-hours and the resulting dataset has shape 4,252,870 *×* 2 *×* 512 *×* 4,096 (variants, var/ref, positions, dimensions) stored as bf16, amounting to ~34 TB.

### Covariance probe

For each sequence direction *s ∈ {* forward, reverse *}*, Evo 2 produces a *K × d* matrix of retained token embeddings, where *K* = 256 and *d* = 4,096. We subtracted the reference embeddings from the alternate embeddings position-wise to obtain **X**_*s*_ *∈* ℝ^*K×d*^. Mean pooling computes *K*^*−*1^ ∑_*i*_ **x**_*s,i*_, a first-order summary that discards how the embedding dimensions co-vary. The covariance probe instead accumulates second-order interactions across the positions.

#### Asymmetric projection

Two affine projections **P**_*s*_, **Q**_*s*_ *∈* ℝ^*d×h*^ (*h* = 64) were learned for each direction and applied to either side of the retained-position covariance, summed over directions and regularized with *ϵ* = 10^*−*3^:

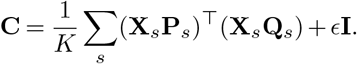

In words, **C** is the truncated covariance under an asymmetric projection. This 64×64 representation captures the most salient pairwise interactions among the projected dimensions and avoids the full 4,096 *×* 4,096 Gram matrix.

#### Spectral regularization

Raw covariance squares the singular values of the embeddings, sharpening the spectrum toward its largest directions, which is undesirable for stable probe training. Hence, we applied a matrix square-root transform to **C**, three iterations of the coupled Newton– Schulz method, which brings the spectrum back to the scale of those singular values. The diagonal *ϵ***I** regularizes the covariance by keeping **C** non-singular.

#### Readout

The regularized covariance 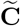 is flattened and mapped by an affine layer to benign and pathogenic logits,

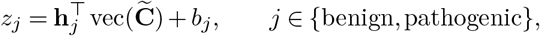

and the pathogenicity prediction is the pathogenic-class softmax probability. The deployed probe has 1,057,026 trainable parameters.

#### Training

The covariance probe was trained with the Muon optimizer (constant learning rate 0.005; batch size 512 *×* 8, mini-batch *×* shards; weight decay 0.1) for a single epoch. Labels were ClinVar P/LP versus B/LB, excluding 0-star. We used an 80/20 gene-level split and did not perform hyperparameter-based model selection. The mean-pooled probe was trained with L2-regularized logistic regression under 5-fold gene-holdout cross-validation, with regularization strength *C* selected via inner 5-fold stratified crossvalidation over 12 values from 10^*−*4^ to 10^1^; features were mean-centered and scaled to unit variance on the training fold. The resulting pathogenicity probe was well-calibrated overall (Supplementary Figure S4).

### Baseline methods

*AlphaMissense*. Pre-computed pathogenicity scores from the AlphaMissense hg38 genome-wide database available for missense variants on MANE Select canonical transcripts only [8].

*CADD*. v1.7 PHRED-scaled scores from pre-computed whole-genome SNV files. Only indels *<*50 bp were scored [15].

*GPN-MSA*. Pre-computed scores were downloaded from HuggingFace (songlab/gpn-msa-hg38-scores) [14].

*NTv3*. Nucleotide Transformer v3 (650M parameters) embeddings were extracted from layer 11 (1,536-D) over 8,192 bp context windows centered on the variant. L2-regularized logistic regression trained on the element-wise difference of alternative and reference embeddings (alt *−* ref) using 5-fold gene-holdout cross-validation [11, 12].

*AlphaGenome*. Variant effect scores were computed using AlphaGenome (google/alphagenome-all-folds) following the published scoring methodology. Reference and alternative gene-body sequences were passed through the model to predict functional genomic tracks across 9 modalities. Scores were aggregated using CenterMask (log_2_ fold-change within variant-centered windows) for chromatin and transcription initiation tracks, and GeneMask (max |ALT *−* REF| or log-fold change over the gene body) for expression and splicing tracks. The final score was the maximum across all modality-specific scores.

### Clinical variant datasets

All benchmarks use ClinVar release 2026-03-21.

#### Full dataset

The full EVEE dataset was built from all of ClinVar by removing multi-allelic, MNV, and unannotated entries (14,118 dropped), then validating against GENCODE v49 for gene-boundary and reference-allele agreement (128,586 dropped), leaving 4,252,870 variants. These divided into a labeled set (1,539,596) and an unlabeled complement (2,716,001). The labeled variants are pathogenic or likely-pathogenic / benign or likely-benign with a non-zero star review status, ensuring at least one submitter provided assertion criteria. In contrast, unlabeled contains all variants of uncertain significance, conflicting interpretations, and zero-star submissions.

#### SNV benchmark

Filtered for SNVs in genes *≤* 100 kb with ≥1 star review status. 833,970 variants with consequence-type breakdown: missense (*n*=125,092, 29.0% pathogenic), synonymous (*n*=292,369, 0.1% pathogenic), nonsense (*n*=36,050, 98.3% pathogenic), splice (*n*=21,683, 98.5% pathogenic), UTR (*n*=31,911, 0.3% pathogenic), intronic (*n*=254,737, 0.7% pathogenic), other (*n*=3,498, 7.4% pathogenic). Gene-level holdout cross-validation ensured no gene appeared in both training and test sets.

#### Indel benchmark

73,961 indels from 3,570 genes *≤*100 kb with *≥*1 star review status. Stratified by consequence: frameshift (*n*=41,398, 99% path.), in-frame (*n*=1,769, 52% path.), noncoding (*n*=24,250, 2% path.), splice-adjacent (*n*=2,442, 88% path.); and by size: 1 bp (*n*=39,801, 70% path.), 2–5 bp (*n*=21,682, 60% path.), 6–20 bp (*n*=9,106, 56% path.), *>*20 bp (*n*=3,372, 68% path.); insertions (*n*=37,558, 61% path.) and deletions (*n*=36,403, 70% path.).

#### Deconfounded benchmark

158,616 variants with pathogenic and benign labels balanced within each consequence type following the framework of Lu et al. [22]. The sampling balances two competing objectives: equalizing pathogenic and benign labels within each consequence type, to prevent models from using consequence as a proxy for pathogenicity, and preserving the natural prevalence of consequence types, to avoid overrepresenting rare categories. This is resolved by weighing each consequence type’s sampling allocation by both its natural prevalence and its amenability to label balance, then selecting variants within each stratum round-robin over genes. This yielded a subset in which consequence-only accuracy fell to 73% from 96% (Supplementary Figure S5).

### Deep mutational scanning datasets

#### BRCA1

Saturation genome editing functional scores from Findlay et al. [23] via MaveDB (urn:mavedb:00000097); 2,803 SNVs spanning 13 exonic regions. Readout: SGE function scores. Variants binarized as functionally abnormal at score *< −*1.22.

#### BRCA2

Saturation genome editing functional classification from Sahu et al. [24] via MaveDB (urn:mavedb:00001225); 6,270 SNVs. Readout: SGE function scores. Variants binarized at score *< −*0.45.

#### TP53

Deep CRISPR mutagenesis functional scores from Funk et al. [25]; 1,920 SNVs across exons 5–8. Readout: Enrich2 relative fitness scores. Variants binarized as functionally abnormal at score *> −*0.97.

#### LDLR

Functional scores measuring cell-surface abundance and LDL uptake from Tabet et al. [26]; 24,999 codon-level variants aggregated to amino acid level by mean pooling. Readout: LDL uptake score. Variants binarized at score *<* 0.40. Target gene fully excluded from ClinVar training for transfer experiments via gene-holdout cross-validation.

### Supervised annotation probes

#### Annotation panel

Annotation probes spanned protein features (disorder content fraction, secondary structure class, backbone psi angle, solvent accessibility, surface area, pLDDT), structural context (chain membership, PPI interface, CpG island, disordered region, Ptc region), regulatory features (histone modification breadth: H3K27me3, H3K4me1, H3K4me3, H3K27ac, overall chromatin breadth), protein domains (Pfam/InterPro families), post-translational modifications (glycosylation, phosphorylation), genomic region (segmental duplication identity, transcript copy number, CDS, intron, replication timing period), amino acid substitution properties (hydrophobicity, volume, molecular weight, Grantham distance, BLOSUM62), and variant effects (splice disruption, charge alteration). Figure 3b reports the 236 binary probes with AUROC measurements.

#### Probe training

For each annotation, a covariance probe was trained on Evo 2 layer 27 embeddings at referencegenome positions to predict the annotation value (the co-variance of a single position is the outer product of the embeddings). Annotations were treated as either binary (classification, evaluated by AUC), categorical (classification, evaluated by accuracy) or continuous (soft binned classification evaluated by Spearman correlation). Training used ~100M positions from~18k genes, with an 80/20 gene-level train/holdout split.

#### Disruption scoring

For each variant, disruption was computed as the change in predicted annotation between alternate and reference sequences. Let *f*_*a*_ be the probe prediction for annotation *a*; the disruption is Δ_*a*_ = *f*_*a*_(**E**_alt_) *f*_*a*_(**E**_ref_).

### Within-gene supervised mechanism-class recovery across four genes

For each of four allelic-heterogeneity genes (LDLR, LMNA, MYH7, TP53), high-confidence ClinVar variants (one- and two-star review status; Pathogenic / Likely Pathogenic) were assigned to literature-curated mechanism classes using a per-gene domain *×* consequence scheme: LDLR *k* = 5 classes (Goldstein–Brown receptor-class taxonomy: null, transport-, binding-, internalization-, and recycling-defective) [32], LMNA *k* = 3 classes (A-band rod DCM/EDMD, R482 hot-spot FPLD2, splice/progeroid) [33], MYH7 *k* = 4 classes (head HCM, converter HCM, mid-rod Laing, distal-rod MSM) [34], TP53 *k* = 4 classes (DNA-contact, structural DBD-destabilizing, null/LoF, tetramerization) [35]. Only variants with a high- or medium-confidence class assignment and class membership *≥* 5 variants were retained, yielding sample sizes of *N* = 1,581 (LDLR), 241 (LMNA), 269 (MYH7), and 488 (TP53).

Per-variant features were the EVEE disruption-profile features (annotation *×* probe Δ scores), filtered (tissue features dropped; |max|*>* 0.05 across the variant set) and standardised per feature. For each gene, a multinomial logistic-regression classifier (L2-regularised, *C* = 0.1) was evaluated by stratified 5-fold cross-validation; balanced accuracy is reported. Baselines evaluated on the same splits were chance (1*/k*), a CADD-only logistic regression on the dbNSFP-derived CADD score, an AlphaMissense-only logistic regression, and a position-only logistic regression on the variant’s encoded residue position. An EVEE + position combination concatenated the positional encoding with the EVEE feature set. Missing per-variant CADD or AlphaMis-sense scores (uncommon at *<* 5% per gene) were mean-imputed within the gene.

### LLM-based interpretation pipeline

For each variant, the top-*k* disruptions (ranked by |Δ|) were assembled alongside variant metadata (gene name, consequence type, genomic position, LOEUF constraint score) into a structured prompt. Claude Sonnet 4.6 [36] was used to synthesize a natural-language interpretation including: a one-sentence summary of the predicted molecular mechanism, a detailed explanation contextualizing the disruption profile against known biology, calibration information (percentage of ClinVar pathogenic variants at similar pathogenicity scores), and bullet-pointed key evidence.

The system prompt (Table S1) describes the disruption signal framework: a probe trained on reference genome activations is run on both reference and variant sequences, producing for each biological feature (helix, conservation, domain membership, etc.) a reference prediction, a variant prediction, and a delta (var *−* ref). The prompt explains that large negative deltas indicate feature disruption while near-zero deltas indicate preservation, and instructs the LLM to lead with the disruption story, use gene function context, and avoid referencing external clinical prediction tools.

The user prompt is assembled programmatically from variant-specific data. It includes: gene name and variant coordinates, consequence type, HGVSp/HGVSc notation, protein domain annotations with LOEUF constraint score, gene function summaries, calibrated pathogenicity percentage, a Region Disruptions table (peak probe signal change across *±*32 kb for the top-10 features by |Δ|*≥* 0.2, with reference, variant, delta, and distance columns), and a Site Disruptions table (probe signal change at the variant position for the top-10 features by |Δ|*≥* 0.1, with reference, variant, and delta columns). An example of the full prompt is shown in Table S2.

### Interpretation evaluation

To evaluate the effect of contextual information on LLM-generated variant interpretations, we conducted a context stratification experiment across 147 expert-reviewed ClinVar variants (*≥*3 stars; 98 benign or likely benign and 49 pathogenic or likely pathogenic). Five cumulative context configurations were tested, each adding information to all previous levels: (1) genomic coordinates only, (2) +gene name, (3) +gene context (domain annotations, LOEUF constraint, gene function summaries), (4) +HGVSp/HGVSc notation, and (5) +Evo 2 disruption profile (calibrated pathogenicity, region and site disruption tables). Configurations without Evo 2 features used a baseline system prompt; the Evo 2 configuration used the probe-aware system prompt described above. Example prompts for each configuration are shown in Table S2.

Interpretations were generated by three Claude model tiers (Haiku 4.5, Sonnet 4.6, and Opus 4.6 [37]). Each interpretation was evaluated by two independent LLM judges (Claude Opus 4.6, Table S3) on three axes using a 1–5 scale: mechanism coverage using a five-level specificity rubric; and biological accuracy and specificity by comparison against expert-submitted ClinVar evidence (SCV records). A composite score was computed as the unweighted mean of the three axes. Confidence intervals represent 95% percentiles. Supplementary Figure S6 shows the composite score stratified by pathogenic and benign variants, and Supplementary Figure S7 shows the per-axis score distributions across context configurations.

#### Example interpretations

Tables S4–S8 show representative LLM interpretations at each mechanism score level (1–5) at the Evo 2 context configuration, illustrating the scoring rubric in practice.

#### Limitations of LLM-as-judge evaluation

Several limitations of this evaluation framework should be noted. First, the ClinVar expert panel evidence used as ground truth varies considerably in mechanistic detail. Some submissions provide extensive molecular reasoning (e.g., the CFTR F508del panel describes misfolding, ER retention, and proteasomal degradation), while others rely primarily on population frequency and computational predictor scores with minimal mechanistic discussion (e.g., the PAH p.Leu213Pro submissions cite REVEL scores and compound heterozygosity but do not describe the structural basis of pathogenicity). This asymmetry means the judge may underestimate mechanism coverage when the interpretation correctly identifies a mechanism that the expert panel simply did not discuss. Second, well-characterized variants in ClinVar (e.g., CFTR F508del, BRCA1 c.5266dupC) are likely present in the LLM’s training data, meaning the interpreting model may draw on memorized knowledge rather than reasoning from the disruption profile alone. The context ablation design partially controls for this, since memorized variants should score well even without Evo 2 features, but subtler forms of memorization cannot be fully ruled out. Third, as with any LLM-as-judge approach, the judge model may have its own biases, and its ability to verify biological claims is limited to what is present in the provided ClinVar evidence.

### RNA-seq validation of EVEE splice and NMD predictions

EVEE-pathogenic variants (score *≥* 0.99, ClinVar *≥* 2 stars) were intersected with CCLE cell-line genotypes (Broad and Sanger WES) and retained when the carrier line provided *≥*5 reads at the variant position and detectable baseline gene expression (CCLE-median junction or transcript count *≥*10). Variants were grouped into three predicted-mechanism tiers: canonical splice donor/acceptor (Tier 1, *±*1*/±*2 from an exon boundary), intronic splice region (Tier 2, +5 / branchpoint), and PTC/frameshift candidates for nonsense-mediated decay (Tier 3). Tier 1/2 variants were called validated if PSI_mut_ *<* 0.30 (*≥*70% of mutant-allele reads use a non-canonical junction); Tier 3 if mutant-allele expression *<* 0.50 *×* CCLE median, VAF-corrected. Multiple cell-line observations per variant were collapsed by taking the most-disrupted observation and the maximum VAF; carriers were classified Hom/LOH (VAF *≥* 0.80) or Het (0.40 *≤* VAF *<* 0.80), with subclonal events (VAF *<* 0.40) excluded. Hom/LOH and Het are reported separately because heterozygous wild-type reads dilute the mutant-allele signal. After aggregation, 82 unique variants remained (Tier 1 *n* = 47; Tier 2 *n* = 3, all Hom/LOH; Tier 3 *n* = 32); validation rates were 30*/*47 (64%) canonical splice, 3*/*3 intronic, 14*/*32 (44%) NMD, and 19*/*23 (83%) Hom/LOH versus 28*/*59 (47%) Het (Figure 4c).

#### Illustrative case

*BRCA1 c*.*5278-1G>A*. BRCA1 c.5278-1G*>*A was inspected in three CCLE ovarian cancer cell lines: JHOS-4 (carrier; VAF 0.96 and 1.00 by Broad and Sanger WES, consistent with germline heterozygosity + somatic loss-of-heterozygosity) and BRCA1 wildtype controls JHOS-2 (sister line) and A2780. Splice-junction reads across BRCA1 exons 19–21 (GENCODE v49, ENST00000357654.9) show zero canonical exon-20 inclusion in JHOS-4, with 24 and 25 reads at the full exon-20 skip junction and the EVEE-predicted cryptic +8 acceptor (8 bp into exon 20), respectively, versus 185–190 (JHOS-2) and 248–294 (A2780) canonical reads in the controls. The 8-bp truncated region (chr17:43,051,109–43,051,116, GRCh38) matches the EVEE-predicted cryptic-acceptor territory.

### Mayo Clinic Tapestry cohort — LDLR clinical-penetrance validation

The Mayo Clinic Tapestry cohort provided per-variant clinical-severity annotations for each LDLR variant carried by a participant. We collapsed the seven-level Tapestry severity annotation into a three-level ordinal scale aligned with standard FH diagnostic rubrics (Dutch Lipid Clinic Network, Simon Broome, MEDPED): *Clinical FH* (definite + probable; both indicating high-confidence FH manifestation), *Suspected FH* (possible; partial diagnostic criteria met), and *Presymptomatic carrier* (LDLR variant present without FH manifestation at ascertainment). Three Tapestry notations were excluded from the severity ordinal as off-axis: hypercholesterolemia of a non-FH lipid pattern (elevated cholesterol not matching the FH pattern, often polygenic or modifier-driven), clinical presentation outside the known FH phenotypic spectrum, and carriers with missing phenotype data.

Tapestry variants were collapsed to one entry per unique variant by protein-level HGVS identifier and intersected with the full LDLR ClinVar EVEE catalogue (4,193 variants), which provides per-variant EVEE scores along with the AlphaMissense and CADD scores from the standard dbNSFP-derived ClinVar annotations. The intersection yielded 147 unique variants (Clinical FH *n* = 111, Suspected FH *n* = 20, Presymptomatic carrier *n* = 16; 83.7% missense, 14.3% truncating or frameshift, 2.0% other). All 147 variants had non-missing scores for every method compared; for non-missense variants in the catalogue, AlphaMissense scores are filled by codon-level imputation (the score of the most pathogenic same-codon missense substitution) and are reported here for comparability. Restricting to the 123 missense variants in the cohort yields AUC= 0.89 for EVEE versus 0.70 for AlphaMissense and 0.61 for CADD. The per-tier distribution of EVEE scores across the three severity tiers is shown in Figure 5b.

Score discrimination was assessed by receiver-operating characteristic AUC for the clinical-screening contrast: positives = Clinical FH *∪* Suspected FH (*n*_+_ = 131), negatives = Presymptomatic carrier (*n*_*−*_ = 16). The Presymptomatic-carrier class is the only true variant-positive / FH-negative comparator available in this cohort and is retained despite class imbalance.

### FinnGen R12 biobank — variant-level *p*-value enrichment

FinnGen R12 GWAS summary statistics [27] (REGENIE; ~500,000 Finnish biobank participants; 2,489 disease endpoints) were obtained from the FinnGen Data Freeze R12 public release. Per-variant EVEE scores and ClinVar classification labels were pre-joined for each variant–phenotype pair across the FinnGen *×* ClinVar overlap. Variants were grouped by EVEE score. The two positive groups were variants with EVEE score *≥* 0.95 (*n* = 958) and the more inclusive *≥* 0.80 set (*n* = 2,497; the former nested within it); the null reference was an allele-frequency–matched control of variants with EVEE score *<* 0.80. As a decircularized check, the *≥* 0.95 group was additionally restricted to variants with no established ClinVar classification (not Pathogenic/Likely pathogenic/Benign/Likely benign; uncertain, conflicting, or unspecified; *n* = 495).

To control for the known confounding effect of allele frequency on GWAS power, the EVEE *<* 0.80 control variants were matched to the allele-frequency distribution of the EVEE *≥* 0.80 group. Minor allele frequency MAF = min(AF, 1 *−* AF) was divided into 30 logarithmically spaced bins spanning [10^*−*5^, 0.5]; within each bin, five control variants were sampled per *≥* 0.80 variant (5:1 matching, with replacement when necessary; seed = 42) to stabilize the null estimate. The control *λ*_GC_ is computed on the full 5:1 set; the control curve is plotted from a subset sized to the *≥* 0.80 group so all three curves span a comparable quantile range.

For each variant, the association *p*-value was taken as the minimum across the 2,489 endpoints and Bonferroni-corrected for the number of endpoints tested. For each group, expected *p*-values were computed as *−* log_10_((rank*−* 0.5)*/N*) after AF matching. The genomic inflation factor *λ*_GC_ was computed as the ratio of the observed median chi-squared statistic to its expected value under the null (0.455). The *y*-axis uses a symmetric log (symlog) scale, switching from linear to *−* log_10_ above *−* log_10_(*p*) = 30.

### ACMG/AMP structured-evidence pipeline (FinnGen R12 variant discovery)

Rare-variant candidates were identified from the FinnGen R12 GWAS summary statistics [27] using a lenient discovery filter: genome-wide significance (*p <* 5 *×* 10^*−*8^) across any binary disease endpoint, minor allele frequency *<* 0.5% in the Finnish population, and an EVEE score *>* 0.8. Variants with active ClinVar Pathogenic / Likely Pathogenic classifications at any star level were excluded, as the pipeline targets variants where reclassification adds value.

Each variant was scored under a structured ACMG/AMP framework incorporating three sequential quality filters and four quantitative criteria. The quality filters applied before scoring were: (i) *bio-coherence*, where the FinnGen phenotype must map to a disease plausibly caused by the implicated gene under its established OMIM condition (e.g. MEPE frameshift + otosclerosis = match; LRBA missense + paraplegia = false positive); (ii) *ClinGen hard filter*, where variants in genes with ClinGen gene–disease validity of “No Known Disease Relationship” or “Contradicted” were excluded, and “Limited” validity blocked PVS1 and PS4 from being applied; (iii) *AR-het flag*, where heterozygous carriers in autosomal-recessive genes cannot satisfy PVS1 (heterozygous LoF is not disease-causing) or PS4 in the standard sense, and were assigned a maximum achievable score of 2 points (PP3 + PM2 only).

The four quantitative criteria were PVS1 (null variant in a gene where loss-of-function is an established disease mechanism), PS4 (case–control odds ratio in FinnGen R12: OR *>* 5 contributes +4 points (*Strong*); OR 2–5, +2 points (*Moderate*); OR 1.5–2, +1 point (*Supporting*)), PP3 (EVEE score *>* 0.5 contributes +1 point, applied at *Supporting* strength as a heuristic pending formal OddsPath calibration under the ClinGen SVI framework [31]; applied to missense candidates only, consistent with PP3’s validated scope), and PM2 (Finnish population allele frequency *<* 0.1% contributes +1 point at *Supporting* strength, per the ClinGen SVI recommendation [38]). Variants were classified by total evidence points under the Tavtigian (2018) point-weighted system [28]: LP-lean (*≥* 4 points), Likely Pathogenic (*≥* 6), and Pathogenic (*≥* 10).

### Mayo Clinic rheumatoid arthritis cohort

We applied EVEE to a clinical exome cohort of 588 patients with confirmed rheumatoid arthritis at Mayo Clinic. Variants were filtered to rare alleles (gnomAD AF *≤* 1%) and annotated against ClinVar, HGMD, REVEL, and CADD, yielding 299 unique rare variants. EVEE scores were obtained for all 299: 257 directly via the public EVEE API and 42 (not yet in the public release) by harvesting Evo 2 7B activations locally and applying the same probe weights, with reproduced scores agreeing with the API to within 5 *×* 10^*−*5^. The 299 variants partition into three ClinVar candidate bins: 225 without an established pathogenic or benign classification (Uncertain significance: 79; Conflicting classifications of pathogenicity: 106; no significance label assigned: 40), 67 likely-benign or background, and 7 known or likely pathogenic. For analysis, variants were stratified by ClinVar candidate bin; high-priority candidates were defined at an EVEE score *≥* 0.90 (14 of the 225 uncertain-classification variants; no benign or likely-benign variant exceeded this threshold). These candidates concentrated in immune-regulation (IFIH1, NOD2, IL10RA) and folate / methotrexate-pathway (ATIC, MTHFR, TYMS) genes, established functional pathways relevant to RA pathophysiology and treatment (Supplementary Figure S11).

## Data availability

Pre-computed pathogenicity predictions and disruption profiles for 4.2 million ClinVar variants are available through the EVEE web application (https://evee.goodfire.ai/) and as a bulk dataset from Zenodo (https://zenodo.org/records/19701997). LLM-based interpretations are generated on demand through EVEE. Programmatic access is provided through the EVEE MCP server (https://github.com/goodfire-ai/evee-mcp).

## Code availability

Computational methods, including an implementation and released weights for covariance-probe inference, cached figure-level data, and scripts to reproduce the main and supplementary figure panels are publicly available via GitHub (https://github.com/goodfire-ai/evee-manuscript).

## Acknowledgements

We thank the Arc Institute and the Evo 2 team for making model weights publicly available, the ClinVar and ClinGen communities for maintaining openly accessible variant interpretation resources, and the participants and investigators of the FinnGen study.

## Author contributions

M.T.P. and N.K.W. conceived the study. T.D. developed the covariance probe methodology. M.T.P., T.D., R.Y., and N.K.W. developed the annotation probe and interpretation framework. S.A., A.J.R. facilitated data extraction and clinical validation in the Mayo Clinic Tapestry cohort. J.M., C.M., S.A., A.J.R., C.O., P.D., B.T., E.M., E.K., P.K., and M.R. provided clinical genomics expertise and variant interpretation guidance. T.K. contributed the FinnGen PheWAS validation. M.B., D.H., and C.F. contributed to model development and evaluation. N.N. and M.A. contributed to infrastructure support for the web app. A.J. and D.B. contributed to project co-ordination and infrastructure. N.K.W. supervised the project. M.T.P., T.D., R.Y., and N.K.W. wrote the manuscript with input from all authors.

---

**Figure S1.**
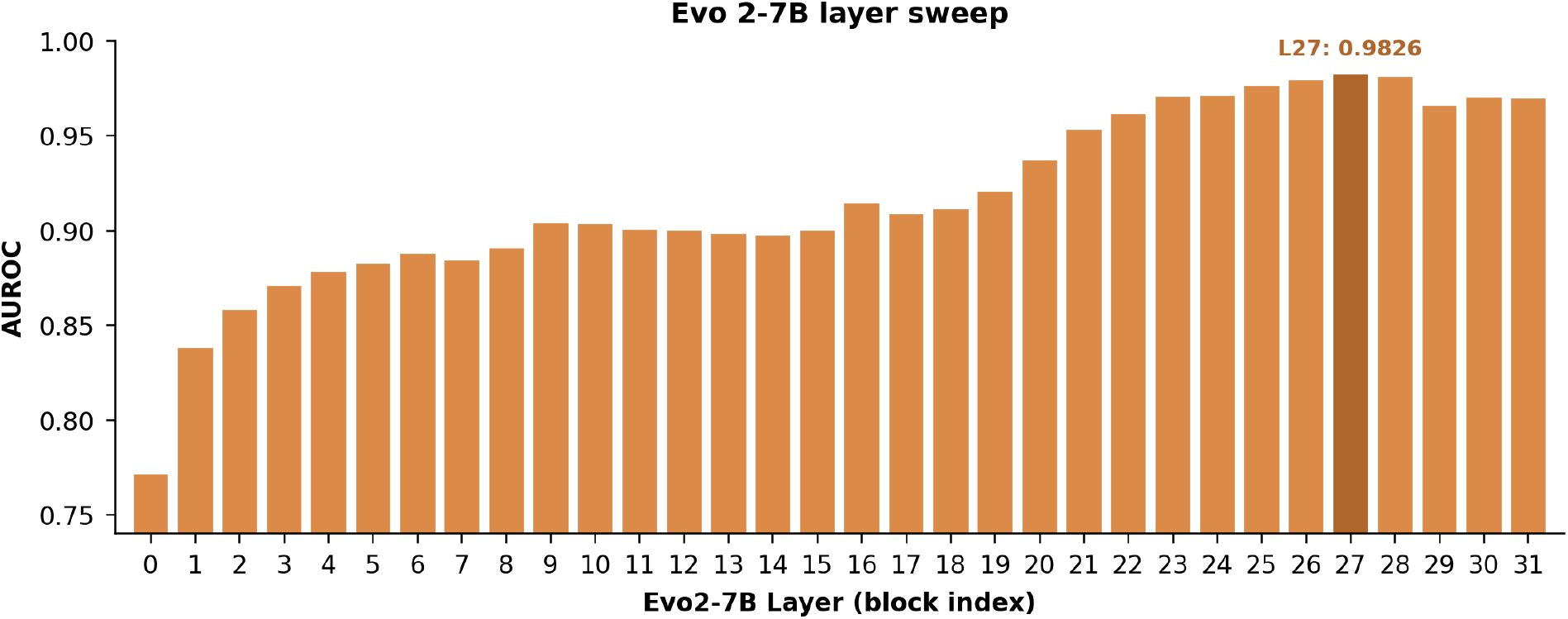
Layer sweep. Layer sweep across all 32 Evo 2-7B blocks showing pathogenicity classification AUROC by layer. Optimal performance at layer 27.

**Figure S2.**
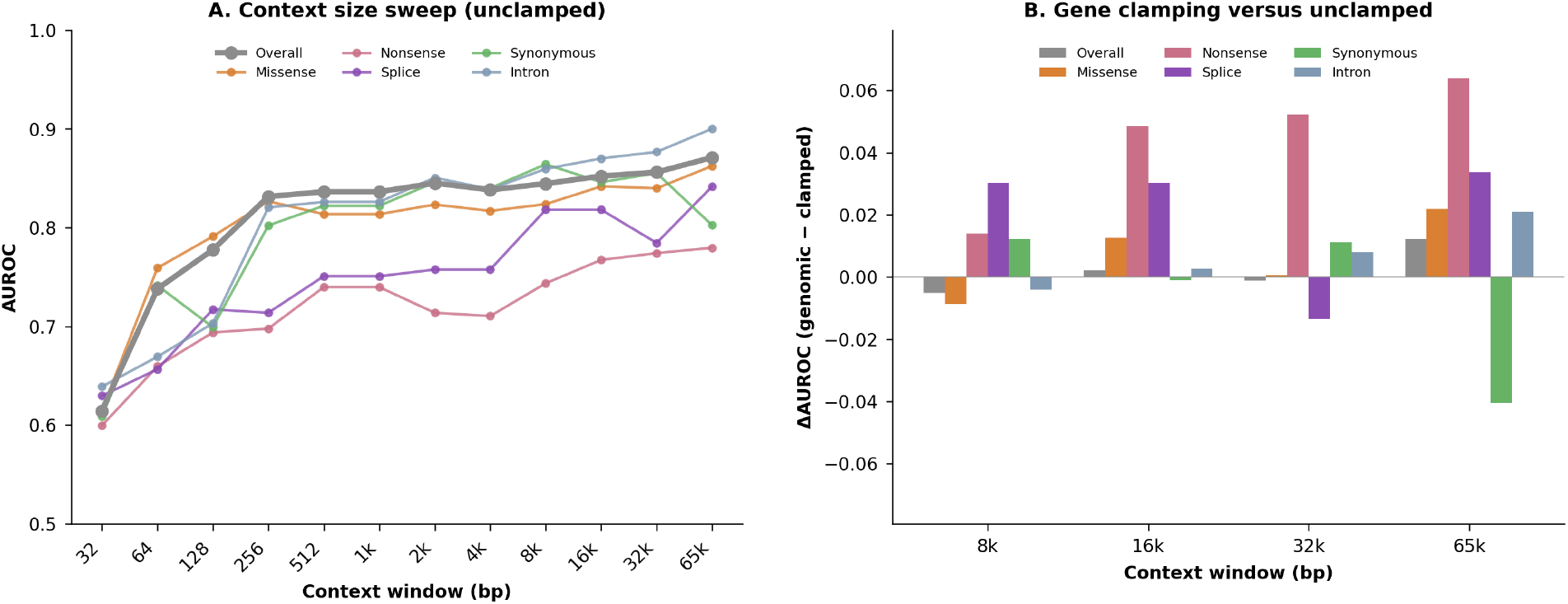
Context window sweep. Context sweeps at layer 27 of Evo 2-7B. Pathogenicity prediction improves sharply with increasing window size below 256 bp, then continues more gradually across all sizes tested. Including flanking sequence beyond gene boundaries outperforms restricting input to the gene body, suggesting Evo 2 captures regulatory signals.

**Figure S3.**
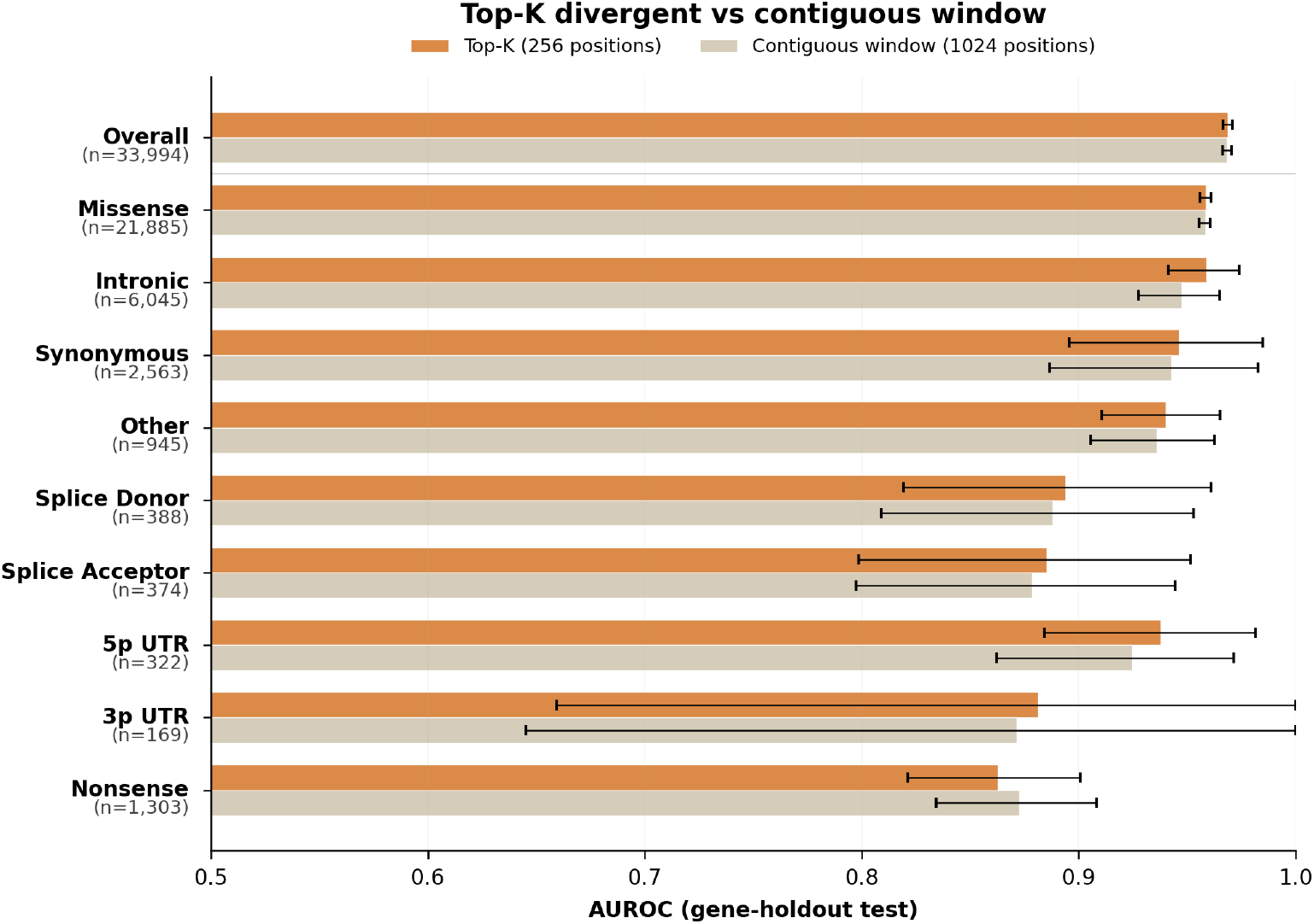
Storage ablation. Comparison of two embedding storage methods: keeping a contiguous window trailing the mutation site or keeping the top-*k* most divergent positions between reference and alternate. Both perform roughly equally well measured by pathogenicity prediction. However, top-*k* better captures far-away secondary effects.

**Figure S4.**
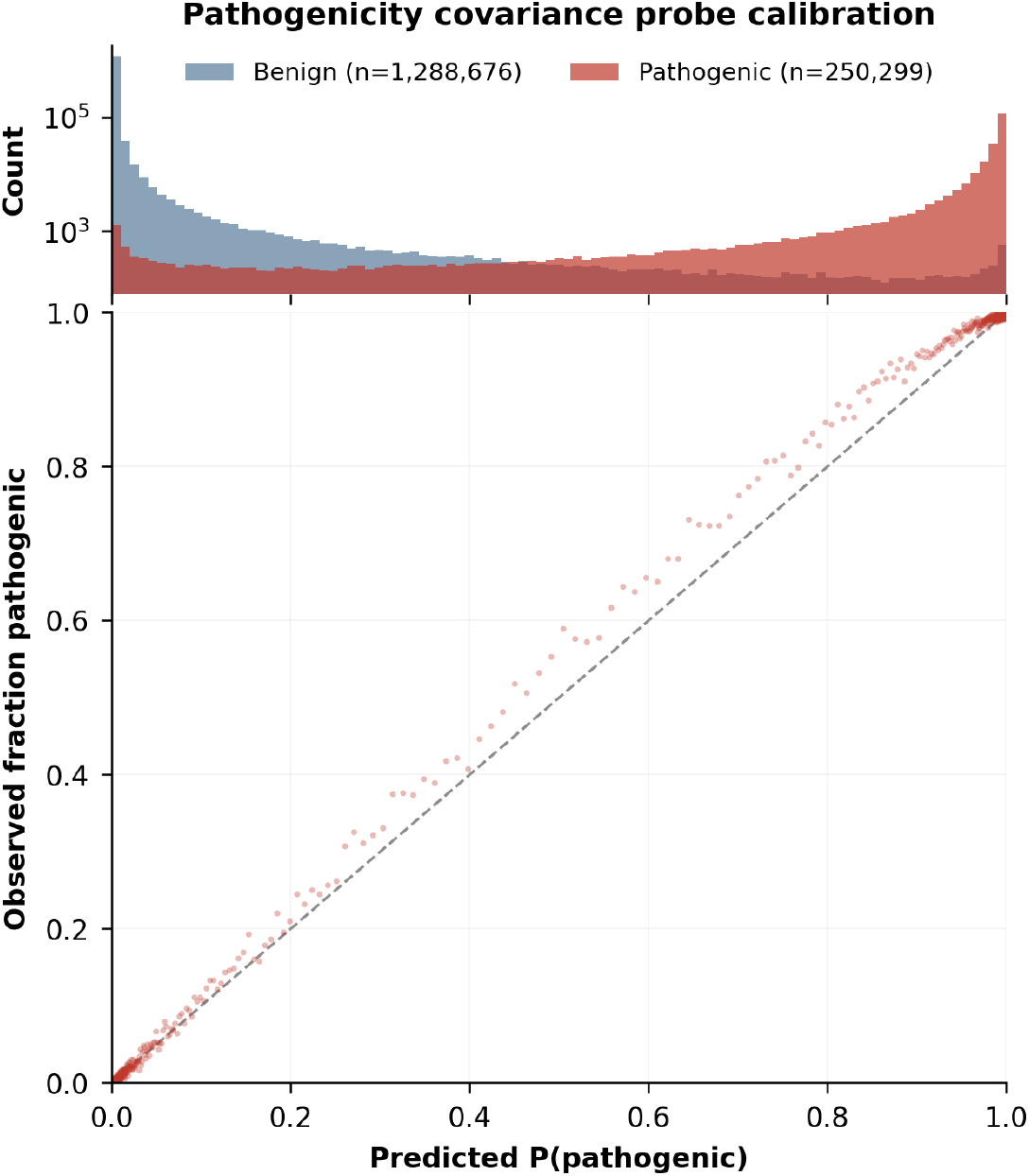
Detailed classification performance. Calibration curve. The pathogenicity probe is well-calibrated overall, especially for confidently benign classifications. It hedges slightly for confidently pathogenic predictions, which is caused by the class imbalance.

**Figure S5.**
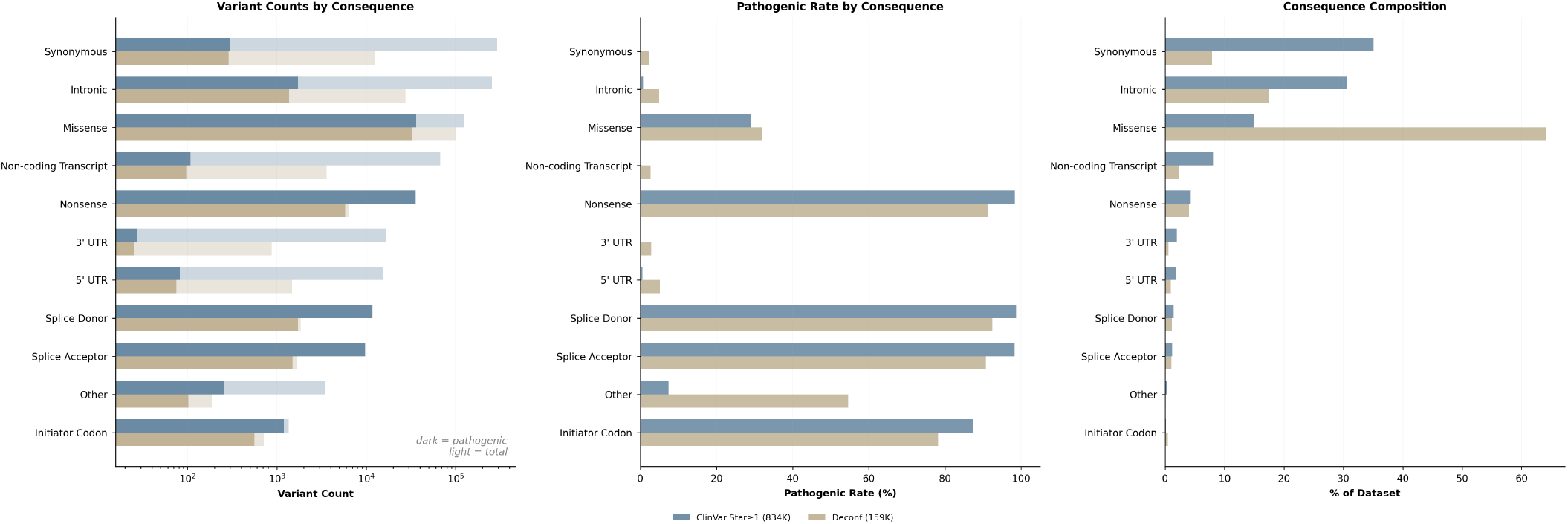
Deconfounded dataset characterization. Variant counts for each consequence type in ClinVar with *≥*1 star (gold) and the deconfounded set (grey) for all variants (light bars) and pathogenic variants (dark bars), showing the downsampling rates. Overall pathogenic rates and composition fraction by consequence in the two datasets.

**Figure S6.**
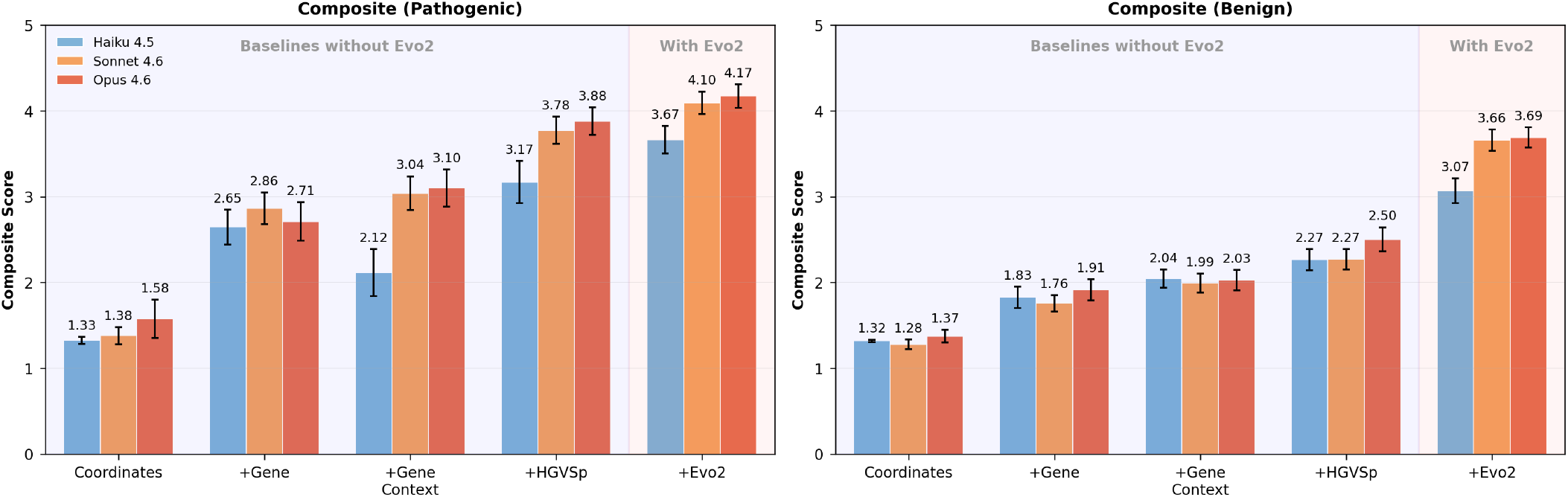
Interpretation quality by pathogenicity class. Composite scores across five context configurations, stratified by pathogenic (left) and benign (right) variants. Three Claude model tiers are compared (Haiku 4.5, Sonnet 4.6, Opus 4.6). Error bars show 95% confidence intervals. Evo 2 predictions provide the largest quality gain for both classes, with pathogenic variants reaching higher absolute scores due to richer mechanistic signals in the disruption profile.

**Figure S7.**
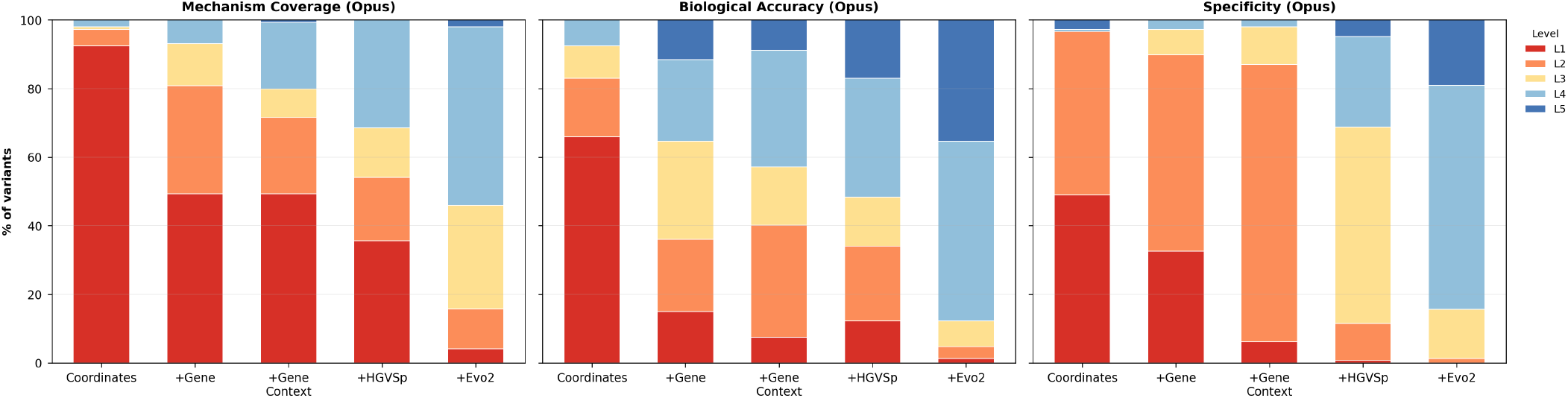
Score distributions across context configurations. Stacked bar charts showing the proportion of variants at each score level (1–5) for mechanism coverage (left), biological accuracy (center), and specificity (right), using Claude Opus 4.6 as the interpreter. Adding Evo 2 predictions shifts the distribution toward higher scores across all three axes, with the most pronounced effect on specificity.

**Figure S8.**
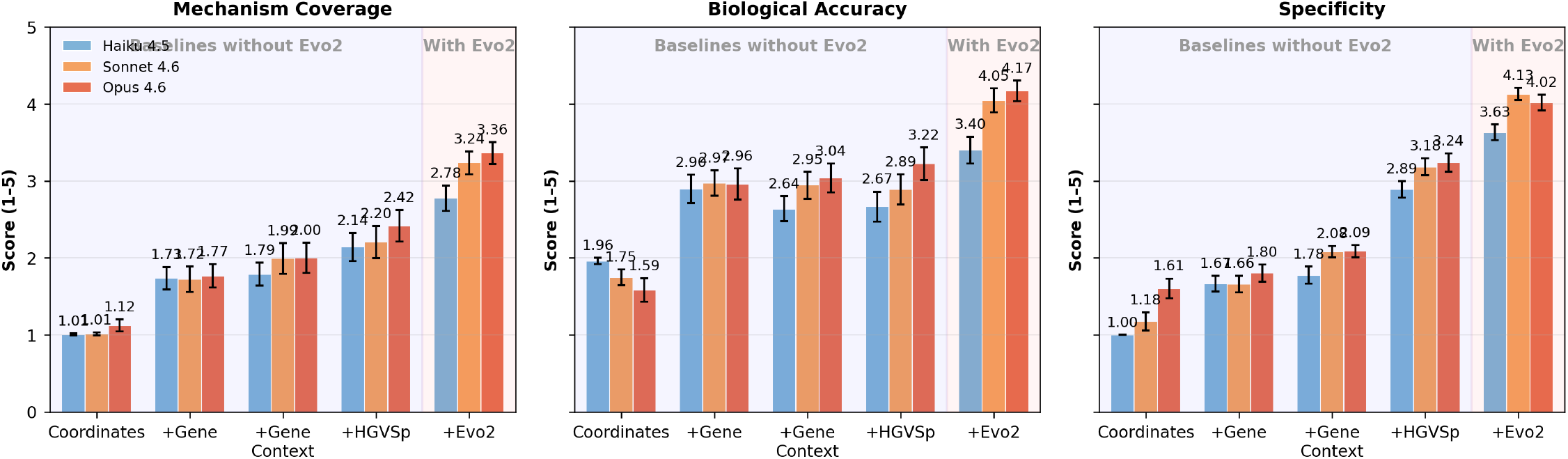
Per-axis interpretation quality across context configurations. Mechanism coverage (left), biological accuracy (center), and specificity (right) scored across the five cumulative context configurations of Figure 3d, for three Claude model tiers. Error bars show 95% confidence intervals. Adding Evo 2 probe predictions improves all three axes, with the largest absolute gain in specificity.

**Figure S9.**
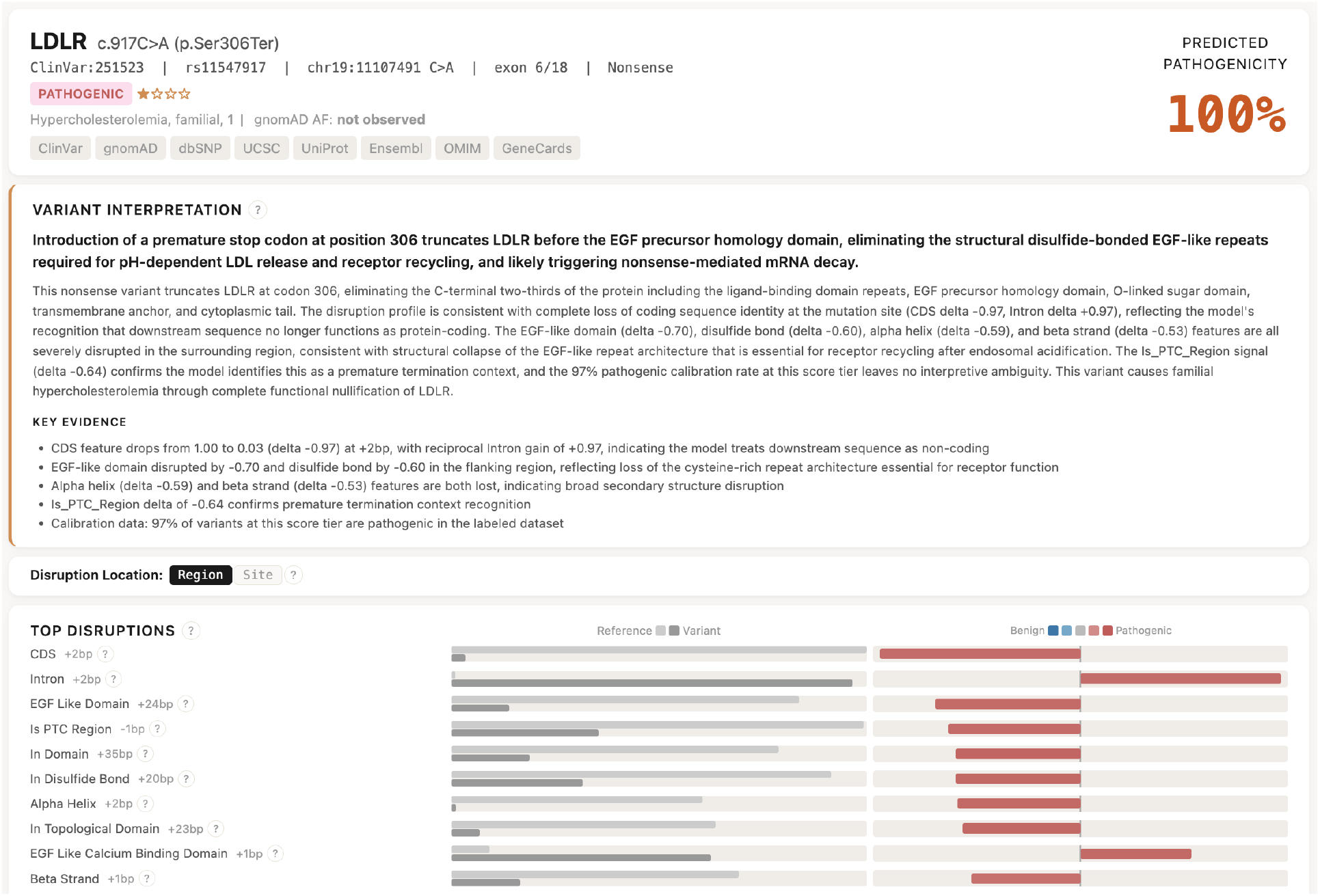
EVEE interpretation for LDLR nonsense variant. LDLR c.917C>A (p.Ser306Ter), predicted 100% pathogenic. The system detects a premature termination codon truncating the protein before the EGF precursor homology domain, with coordinated loss of EGF-like domain (Δ = *−*0.70), disulfide bond (Δ = *−*0.60), and alpha helix (Δ = *−*0.89) features. The interpretation correctly links these disruptions to loss of the receptor recycling machinery required for LDL uptake.

**Figure S10.**
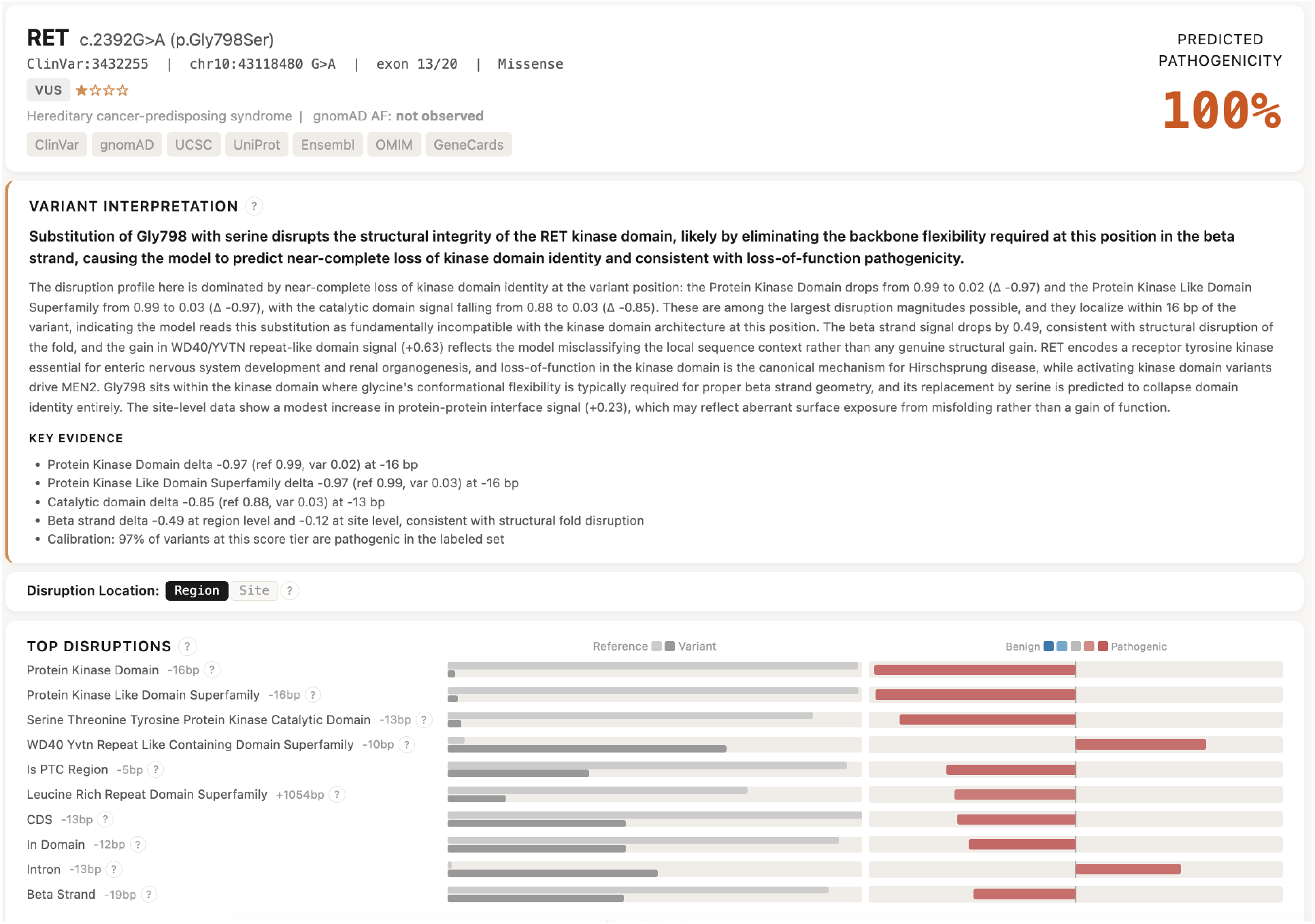
EVEE interpretation for RET missense variant. RET c.2392G>A (p.Gly798Ser), predicted 100% pathogenic. The system detects near-complete loss of protein kinase domain (Δ = *−*0.97) and catalytic domain (Δ = *−*0.65) annotations with concurrent beta strand disruption (Δ = *−*0.48). The interpretation correctly identifies that glycine’s backbone flexibility is required at this kinase fold position, and serine substitution causes structural disruption consistent with loss of function.

**Figure S11.**
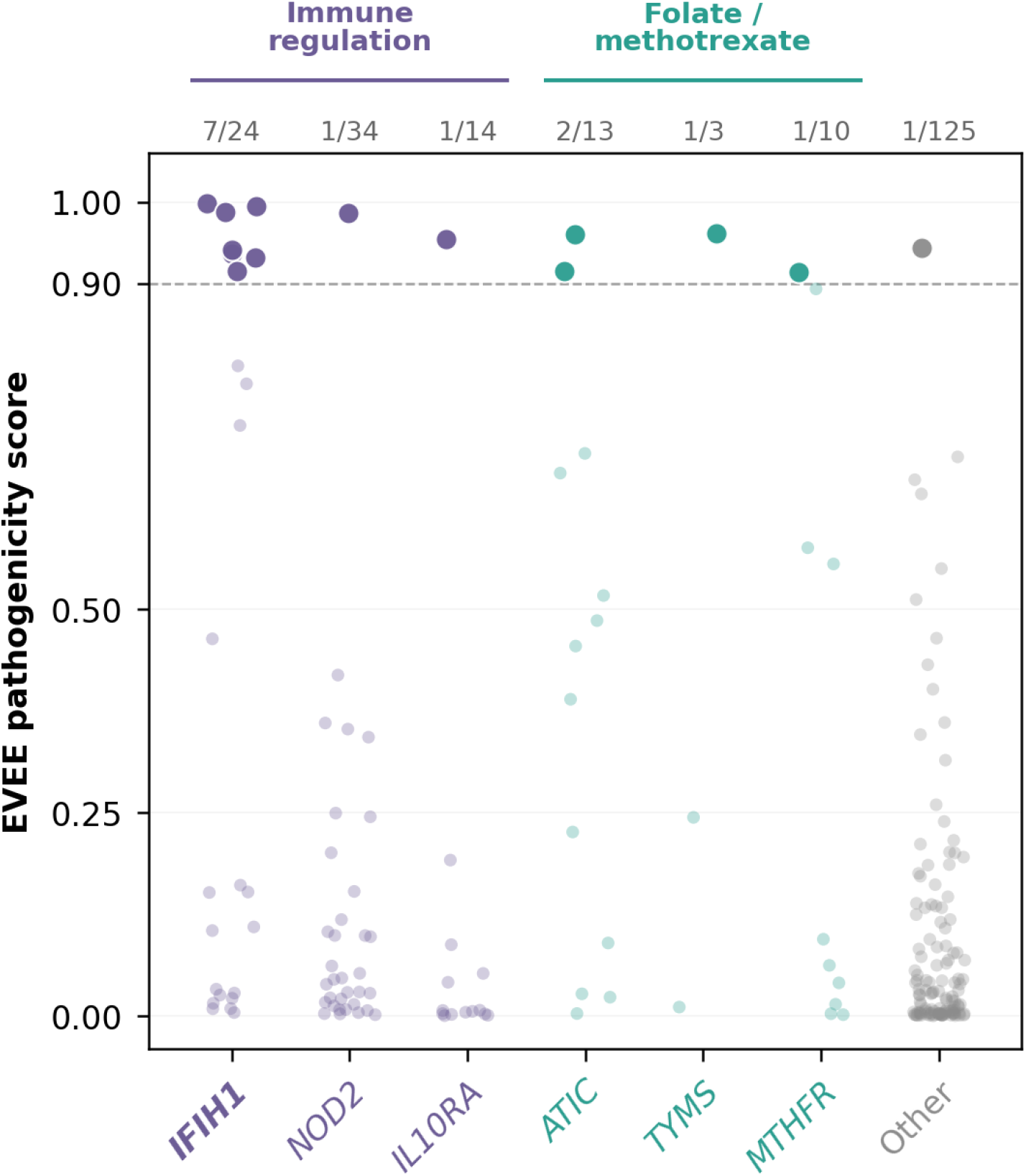
Clinical-deployment demonstration in a Mayo Clinic rheumatoid arthritis cohort. Gene-level EVEE scores for the 225 of 299 cohort rare variants (gnomAD AF *≤* 1%) lacking an established pathogenic or benign ClinVar classification (uncertain, conflicting, or unspecified significance), drawn from 588 patients with confirmed rheumatoid arthritis at Mayo Clinic. Saturated dots score *≥* 0.90 (14 variants; no benign or likely-benign cohort variant exceeds this threshold); high-priority candidates concentrate in immune regulation (IFIH1, NOD2, IL10RA; purple) and the folate / methotrexate pathway (ATIC, MTHFR, TYMS; teal). Per-column tallies show the high-scoring count over total cohort variants per gene. The pipeline compresses 299 rare candidate variants into a mechanism-annotated shortlist suitable for expert clinical review.

**Table S1.** System prompts for variant interpretation. Two system prompts were used depending on whether the context configuration includes Evo 2 probe features. Both prompts request a structured JSON response containing a 3–5 sentence summary, a one-sentence mechanism, a confidence level (high/moderate/low), and key evidence points. Both include identical writing-style constraints: direct, concise prose; no em dashes; no superlatives or hedging language; no repetition of the variant ID; specific numbers preferred over vague qualifiers.

| Prompt | Content |
| --- | --- |
| Baseline<br>(without Evo2) | You are a clinical genomics expert. Based on the variant information provided, interpret the likely biological impact and pathogenicity of this variant.<br>Be specific about biological mechanisms when the information supports it. 3–5 sentences for summary, 1 sentence for mechanism.<br>Writing style: Write like a senior geneticist. Be direct and concise. Never use em dashes. Never use “notably”, “critically”, “importantly”, “striking”. Don’t use superlatives or hedging language. State facts plainly. Don’t repeat the variant ID or coordinates in the summary. Specific numbers are good. Vague qualifiers (“profound”, “massive”, “broad”) are not.<br>Return JSON: {summary, mechanism, confidence, key_evidence} |
| Evo2<br>(with Evo2) | You are a clinical genomics expert interpreting variant pathogenicity predictions from Evo2, a 7-billion-parameter DNA foundation model.<br>The system uses a probe trained on reference genome activations, then run on both reference and variant activations. For each biological feature (helix, conservation, domain membership, etc.), you see: **ref** (the probe’s prediction on the reference genome), **var** (the prediction on the variant genome), and $\Delta$ (var – ref, the disruption signal). Large negative deltas mean the variant disrupts that feature. Near-zero deltas mean the feature is preserved. This is the key signal: pathogenic variants show large disruptions, benign variants show near-zero deltas.<br>Provide a clinical interpretation: lead with the disruption story (which features were disrupted and which preserved); use the gene function context to explain why disruption of those features matters for this gene; explain why the model predicts what it does based on the disruption profile. Do not reference clinical prediction tools (CADD, AlphaMissense, SIFT, PolyPhen, etc.). 3–5 sentences for summary, 1 sentence for mechanism.<br>Writing style: Write like a senior geneticist. Be direct and concise. Never use em dashes. Never use “notably”, “critically”, “importantly”, “striking”. Don’t use superlatives or hedging language. State facts plainly. Don’t repeat the variant ID or coordinates in the summary. Specific numbers are good. Vague qualifiers (“profound”, “massive”, “broad”) are not.<br>Return JSON: {summary, mechanism, confidence, key_evidence} |

**Table S2.**
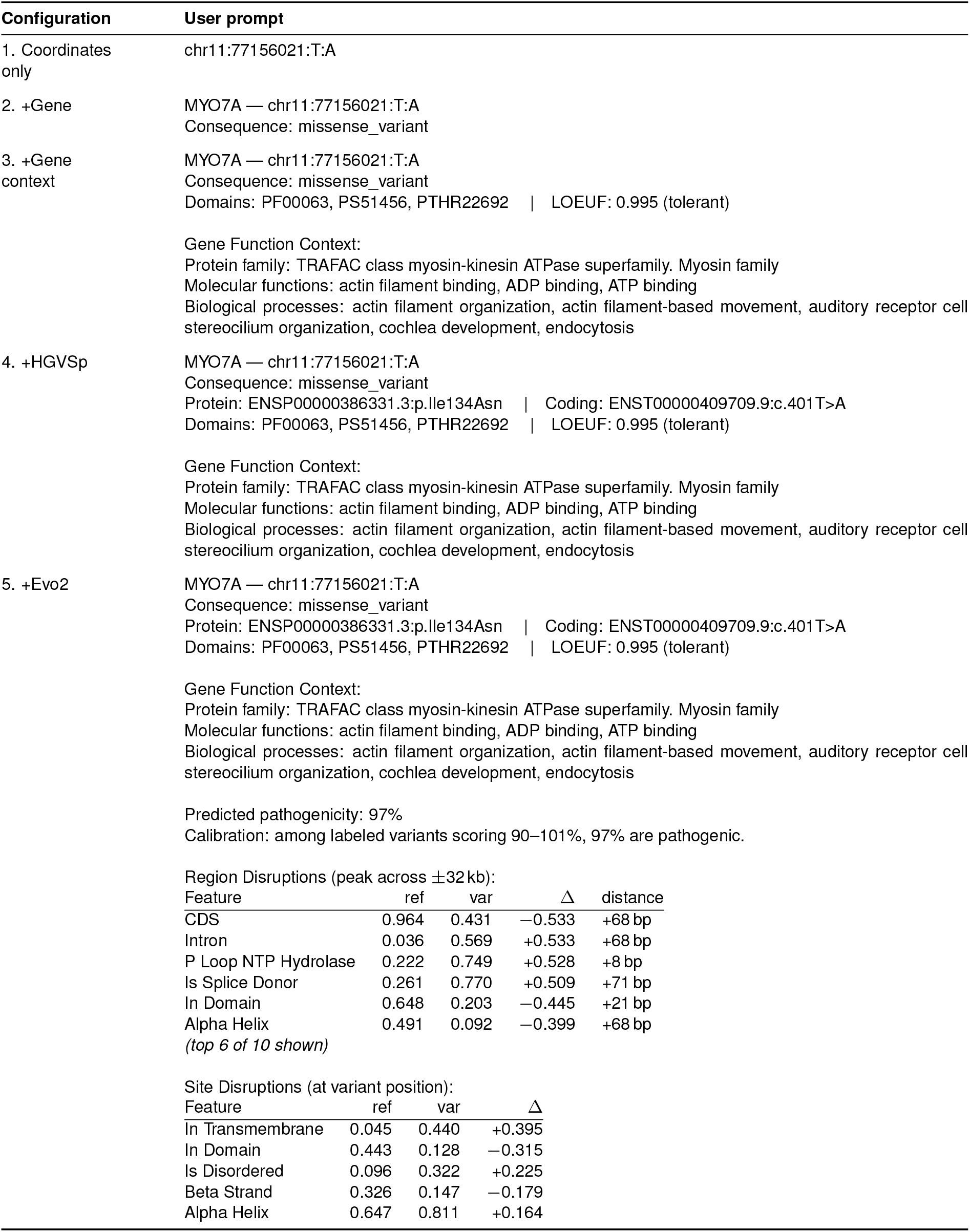
Example user prompts for each context configuration. Each prompt shows the same variant (MYO7A chr11:77156021:T:A, a pathogenic missense variant) at increasing context levels. Each configuration cumulatively adds information to the previous level. The Evo 2 configuration uses the Evo 2 system prompt (Table S1); all others use the baseline system prompt.

**Table S3.** LLM judge prompts. Two independent judges (Claude Opus 4.6) evaluate each interpretation. Each judge receives the variant identifier, the LLM-generated interpretation, and ClinVar expert panel evidence as context.

| Judge | Prompt |
| --- | --- |
| Mechanism coverage | You are evaluating an automated variant interpretation generated by an AI system that uses probe features from a genomic language model (Evo2) and an LLM without internet access. The system has no access to patient data, population frequencies, functional assays, or experimental results. Use the ClinVar expert panel evidence as ground truth for the molecular mechanism; ignore any non-expert submissions.<br>*Variant:* {variant_id}<br>*LLM Interpretation:* {interpretation}<br>*ClinVar Expert Panel Evidence (ground truth):* {clinvar_evidence}<br>*Mechanism Coverage Rubric (1–5).* Score = highest specificity level of mechanism correctly identified, using ClinVar expert panel SCV as ground truth. Only correct claims count.<br>Level 1: **Gene-level only.** Names gene/disease/pathway but no variant-specific mechanism.<br>Level 2: **Correct variant-type.** Identifies consequence type and expected effect but not which domain/residue.<br>Level 3: **Correct functional class or protein region.** Names the right domain or functional category.<br>Level 4: **Specific mechanism with biological reasoning.** Connects variant to consequence through biological logic.<br>Level 5: **Level 4 + additional biological detail.** Second affected function, biophysical basis, downstream pathway, or non-obvious mechanism.<br>*Judge procedure:* (1) Extract mechanism claims from the expert SCV; (2) extract mechanism claims from the LLM interpretation; (3) compare, where correct claims match or are consistent with the SCV and incorrect claims contradict it; (4) assign the score as the highest rubric level among correct claims.<br>*Scoring rules:* Wrong pathogenicity direction caps the score at 1; hedged claims score at the level described if correct; missing clinical, population, or experimental data is not penalized.<br>Returns JSON: {mechanism_score: 1–5, reasoning: 2–3 sentence justification}. |
| Biological accuracy & specificity | You are evaluating an automated variant interpretation generated by an AI system that uses probe features from a genomic language model (Evo2).<br>*Variant:* {variant_id}<br>*LLM Interpretation:* {interpretation}<br>*ClinVar Submitted Evidence (human-curated):* {clinvar_evidence}<br>Compare the LLM interpretation against the ClinVar evidence. Score on two axes (1–5):<br>(1) *biological_accuracy:* Are the biological claims in the interpretation consistent with the ClinVar evidence? (5 = fully consistent, 1 = contradicts evidence)<br>(2) *specificity:* Does the interpretation make specific, concrete claims rather than generic statements? (5 = very specific to this variant, 1 = could apply to any variant)<br>Returns JSON: {biological_accuracy: 1–5, specificity: 1–5}. |

**Table S4.** Example interpretation: mechanism score 1 (gene-level only). *ETHE1* c.55G>T (chr19:43506794:C:A), classified as benign by the ClinGen Mito Variant Curation Expert Panel, generated at the Evo 2 context configuration by Claude Opus 4.6. The LLM misidentifies the coding variant as a 3’ UTR variant (c.*55G>T), building its entire mechanism reasoning on an incorrect premise. While the benign conclusion is correct, only gene-level identification is accurate.

| Component | Content |
| --- | --- |
| LLM summary | This 3' UTR variant in *ETHE1* shows a 0% pathogenicity prediction, consistent with the absence of any meaningful disruption to protein-coding or structural features. As a variant located 55 bases past the stop codon, it falls outside the translated region and is not expected to alter protein structure, active site residues, or conserved functional domains. The LOEUF score of 0.979 indicates *ETHE1* is relatively tolerant to loss-of-function variation, further reducing concern. The model correctly identifies this as a benign variant given the lack of disruption to any feature relevant to *ETHE1*'s sulfur dioxygenase or iron-binding functions. |
| LLM mechanism | A 3' UTR variant at this position does not disrupt the coding sequence, protein folding, or any catalytic domain required for *ETHE1*'s mitochondrial sulfur dioxygenase activity. |
| Expert SCV | The filtering allele frequency of the c.55G>T variant in the *ETHE1* gene is >0.7% by the Exome Aggregation Consortium, which is a high enough frequency to be classified as benign based on thresholds defined by the ClinGen Mito Variant Curation Expert Panel (BA1). |
| Judge reasoning | The LLM correctly identifies the gene (*ETHE1*) and reaches the correct benign conclusion, but incorrectly classifies the variant as a 3' UTR variant (c.*55G>T) when the expert panel identifies it as c.55G>T (coding region). Since the variant-type mechanism reasoning is based on an incorrect premise about the variant's location, the highest correct claim is gene-level identification with correct pathogenicity direction. |

**Table S5.** Example interpretation: mechanism score 2 (correct variant-type). *MAP2K1* c.1068+9A>G (chr15:66489771:A:G), classified as likely benign by the ClinGen RASopathy Expert Panel, generated at the Evo 2 context configuration by Claude Opus 4.6. The LLM correctly identifies the intronic variant as outside the canonical splice site with no predicted disruption, but does not advance beyond varianttype classification.

| Component | Content |
| --- | --- |
| LLM summary | This intronic variant located 9 bases into the intron shows no disruptive signal across any probed feature. All deltas are near zero or slightly positive, with no negative disruptions to secondary structure, domain integrity, or protein interactions. The small positive shifts in kinase domain and secondary structure probes at a nearby position likely reflect minor probe noise rather than meaningful biological change, as the variant sits well outside the critical splice donor dinucleotide. *MAP2K1* is a constrained gene (LOEUF 0.515), but this deep intronic position is unlikely to affect splicing or protein function. The model assigns 0% pathogenicity, consistent with a benign intronic variant that preserves all functional features. |
| LLM mechanism | An intronic variant at position c.1068+9 that does not disrupt splice signals, protein domain structure, or any other probed biological feature. |
| Expert SCV | The c.1068+9A>G intronic variant in the *MAP2K1* gene is classified as likely benign because it does not alter an amino acid residue, is not located within the canonical splice site, and computational splice prediction tools do not predict an impact on splicing (BP4, BP7). It has been identified in 0.01123% of Latino chromosomes in gnomAD (BS1 not met). In summary, this variant meets criteria to be classified as likely benign. ACMG/AMP criteria applied: BP4, BP7. |
| Judge reasoning | The LLM correctly identifies this as an intronic variant (c.1068+9) outside the canonical splice site with no predicted splice disruption, matching the expert SCV's BP4/BP7 classification as likely benign. This represents correct variant-type identification (intronic, no splice impact) consistent with Level 2. While the LLM mentions the kinase domain and specific distance from the splice site, these details support the same "intronic, no splice impact" conclusion rather than adding a distinct functional region or mechanistic insight that would reach Level 3. |

**Table S6.**
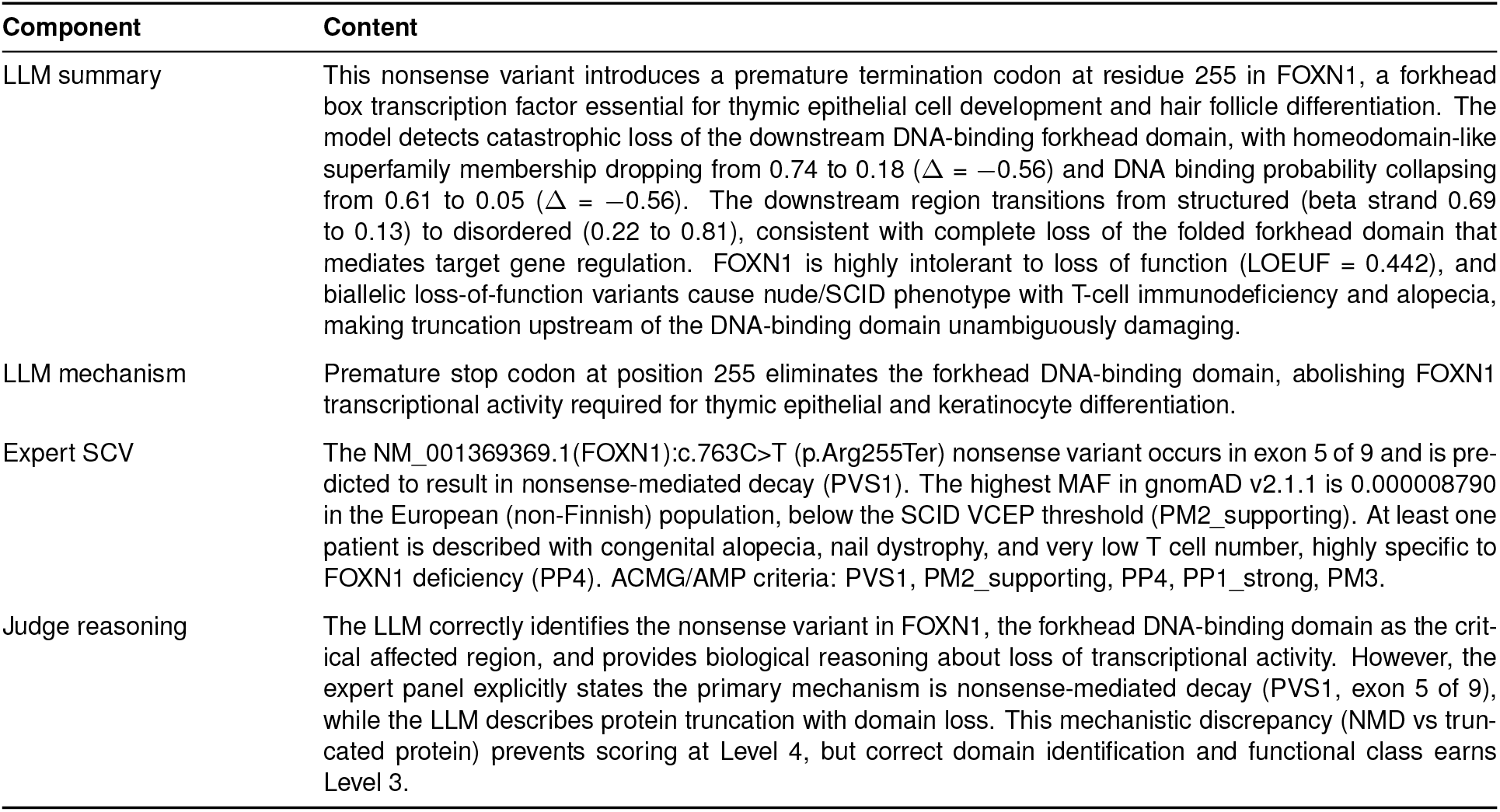
Example interpretation: mechanism score 3 (correct functional class). *FOXN1* p.Arg255Ter (chr17:28529156:C:T), classified as pathogenic by the ClinGen SCID VCEP, generated at the Evo 2 context configuration by Claude Opus 4.6. The LLM correctly localizes the nonsense variant to the forkhead DNA-binding domain, but proposes protein truncation with domain loss rather than the expert-annotated nonsense-mediated decay mechanism.

**Table S7.** Example interpretation: mechanism score 4 (specific mechanism with biological reasoning). *PAH* p.Leu213Pro (chr12:102855203:A:G), classified as pathogenic by multiple clinical laboratories, generated at the Evo 2 context configuration by Claude Opus 4.6. The LLM connects proline-mediated helix breaking to destabilization of the catalytic domain required for enzymatic activity.

| Component | Content |
| --- | --- |
| LLM summary | This Leu-to-Pro substitution in the PAH catalytic domain produces a clear alpha helix disruption at the variant site ($\Delta$ $-0.245$, from $0.975$ to $0.731$), with a concurrent gain in beta strand propensity ($+0.218$) and increased disorder ($+0.166$). Proline is a well-known helix breaker, and the probe captures exactly this structural consequence: a position that was confidently helical in the reference is shifted toward irregular or strand-like conformation. The variant also increases the predicted protein-protein interaction interface signal ($+0.147$), suggesting altered surface properties at a region involved in PAH oligomerization or cofactor interaction. PAH requires precise active-site geometry for iron coordination and BH4 cofactor binding, and destabilization of a helix within the catalytic domain (PF00351) is a recognized mechanism for loss of phenylalanine hydroxylase activity, consistent with classical PKU. |
| LLM mechanism | Proline substitution disrupts an alpha helix in the PAH catalytic domain, increasing local disorder and destabilizing the structural framework required for iron binding and enzymatic activity. |
| Expert SCV | This sequence change replaces leucine, which is neutral and non-polar, with proline, which is neutral and non-polar, at codon 213 of the PAH protein (p.Leu213Pro). This variant is not present in population databases (gnomAD no frequency). This missense change has been observed in individuals with phenylketonuria (PMIDs: 8632937, 9521426, 19292873, 22112818, 23430547). Advanced modeling indicates that this missense variant is expected to disrupt PAH protein function with a positive predictive value of 80%. PAH-specific ACMG/AMP criteria applied: PP4_Moderate, PM3_Very-strong. |
| Judge reasoning | The LLM correctly identifies the pathogenic direction, places the variant in the PAH catalytic/hydroxylase domain (PF00351), and provides specific biological reasoning connecting proline-mediated helix disruption to destabilization of the structural framework required for iron binding and enzymatic activity. This is consistent with the expert panel's pathogenic classification and PP3 computational evidence, reaching Level 4 specificity with mechanistic biological logic beyond just naming the domain. |

**Table S8.** Example interpretation: mechanism score 5 (multi-mechanism with biophysical detail). *GAA* p.Met1Val (chr17:80104586:A:G), classified as pathogenic by the ClinGen Lysosomal Diseases VCEP for Pompe disease, generated at the Evo 2 context configuration by Claude Opus 4.6. The LLM traces a multi-step biological chain from start codon loss through signal peptide elimination to failed lysosomal targeting, entirely from disruption profile signals.

| Component | Content |
| --- | --- |
| LLM summary | This variant abolishes the initiator methionine codon of GAA, with the model detecting a start codon probability drop from $0.976$ to $0.477$ at the variant site and from $0.959$ to $0.079$ in the peak region ($+2$ bp). The downstream consequences are severe: the model predicts loss of the signal/transit peptide ($0.382$ to $0.029$), loss of the protein chain ($0.863$ to $0.230$), and near-complete destruction of the transmembrane signal peptide region ($0.923$ to $0.007$ at $+48$ bp). Without a functional start codon, GAA acid alpha-glucosidase cannot be synthesized or properly targeted to lysosomes, which is the direct cause of Pompe disease (glycogen storage disease type II). The model also detects aberrant CDS and polypyrimidine tract signals upstream, consistent with the translational machinery failing to recognize this region as a normal coding sequence initiation site. |
| LLM mechanism | Loss of the initiator methionine codon prevents translation of GAA acid alpha-glucosidase, eliminating the signal peptide required for lysosomal targeting and abolishing enzymatic glycogen degradation. |
| Expert SCV | The NM_000152.5:c.1A>G (p.Met1Val) start loss variant in GAA may cause a truncated or absent protein by altering the start codon (PVS1_Strong). This variant is absent in gnomAD v2.1.1 (PM2_Supporting). This variant has been detected in at least 5 individuals with Pompe disease, including three compound heterozygous for another pathogenic variant (PM3_Strong). At least two patients had documented GAA deficiency with $<10\%$ of the normal mean control level of GAA activity in dried blood spots. One homozygous patient had documented severe infantile onset and was CRIM negative (PP4_Moderate). ACMG/AMP criteria: PVS1_Strong, PM3_Strong, PP4_Moderate, PM2_Supporting. |
| Judge reasoning | The LLM correctly identifies the start codon loss in GAA causing absent or truncated protein, matching the SCV's p.Met1Val start loss, and goes beyond by explaining the biological chain: loss of initiator methionine, loss of signal peptide, failure of lysosomal targeting, abolished glycogen degradation, and Pompe disease. This provides Level 4 biological reasoning (connecting variant to consequence) plus additional biological detail about the signal peptide and lysosomal targeting pathway, reaching Level 5. |

## References

1. Zeming Lin, Halil Akin, Roshan Rao, Brian Hie, Zhongkai Zhu, Wenting Lu, Nikita Smetanin, Robert Verkuil, Ori Kabeli, Yaniv Shmueli, Allan dos Santos Costa, Maryam Fazel-Zarandi, Tom Sercu, Salvatore Candido, and Alexander Rives. Evolutionary-scale prediction of atomic-level protein structure with a language model. Science, 379(6637):1123– 1130, 2023. doi: 10.1126/science.ade2574.

2. Garyk Brixi, Matthew G Durrant, Jerome Ku, Mohsen Naghipourfar, Michael Poli, et al. Genome modelling and design across all domains of life with Evo 2. Nature, 652: 1349–1361, 2026. doi: 10.1038/s41586-026-10176-5.

3. Konrad J Karczewski, Laurent C Francioli, Grace Tiao, Beryl B Cummings, Jessica Alföldi, Qingbo Wang, Ryan L Collins, Kristen M Laricchia, Andrea Ganna, Daniel P Birnbaum, et al. The mutational constraint spectrum quantified from variation in 141,456 humans. Nature, 581(7809):434–443, 2020. doi: 10.1038/s41586-020-2308-7.

4. Lea M Starita, Nadav Ahituv, Maitreya J Dunham, Jacob O Kitzman, Frederick P Roth, Georg Seelig, Jay Shendure, and Douglas M Fowler. Variant interpretation: functional assays to the rescue. The American Journal of Human Genetics, 101 (3):315–325, 2017. doi: 10.1016/j.ajhg.2017.07.014.

5. Melissa J Landrum, Jennifer M Lee, Mark Benson, Garth Brown, Chen Chao, Shanmuga Chitipiralla, Baoshan Gu, Jennifer Hart, Douglas Hoffman, Jeffrey Hoover, et al. Clinvar: public archive of interpretations of clinically relevant variants. Nucleic Acids Research, 44(D1):D862–D868, 2016. doi: 10.1093/nar/gkv1222.

6. Sue Richards, Nazneen Aziz, Sherri Bale, David Bick, Soma Das, Julie Gastier-Foster, Wayne W Grody, Madhuri Hegde, Elaine Lyon, Elaine Spector, et al. Standards and guidelines for the interpretation of sequence variants: a joint consensus recommendation of the American College of Medical Genetics and Genomics and the Association for Molecular Pathology. Genetics in Medicine, 17(5):405–424, 2015. doi: 10.1038/gim.2015.30.

7. Jonathan Frazer, Pascal Notin, Mafalda Dias, Aidan Gomez, Joseph K Min, Kelly Brock, Yarin Gal, and Debora S Marks. Disease variant prediction with deep generative models of evolutionary data. Nature, 599(7883):91–95, 2021. doi: 10.1038/s41586-021-04043-8.

8. Jun Cheng, Guido Novati, Joshua Pan, Clare Bycroft, Akvilė Žemgulytė, Taylor Applebaum, Alexander Pritzel, Lai Hong Wong, Michal Zielinski, Tobias Sargeant, et al. Accurate proteome-wide missense variant effect prediction with AlphaMissense. Science, 381(6664):eadg7492, 2023. doi: 10.1126/science.adg7492.

9. Žiga Avsec, Vikram Agarwal, Daniel Visentin, Joseph R Ledsam, Agnieszka Grabska-Barwinska, Kyle R Taylor, Yannis Assael, John Jumper, Pushmeet Kohli, and David R Kelley. Effective gene expression prediction from sequence by integrating long-range interactions. Nature Methods, 18(10): 1196–1203, 2021. doi: 10.1038/s41592-021-01252-x.

10. Johannes Linder, Divyanshi Srivastava, Han Yuan, Vikram Agarwal, and David R Kelley. Predicting RNA-seq coverage from DNA sequence as a unifying model of gene regulation. Nature Genetics, 57:949–961, 2025. doi: 10.1038/s41588-024-02053-6.

11. Sam Boshar, Benjamin Evans, Ziqi Tang, Armand Picard, Ya-nis Adel, Franziska K. Lorbeer, Chandana Rajesh, Tristan Karch, Shawn Sidbon, David Emms, Javier Mendoza-Revilla, Fatimah Al-Ani, Evan Seitz, Yair Schiff, Yohan Bornachot, Ariana Hernandez, Marie Lopez, Alexandre Laterre, Karim Beguir, Peter Koo, Volodymyr Kuleshov, Alexander Stark, Bernardo P. de Almeida, and Thomas Pierrot. A foundational model for joint sequence-function multi-species modeling at scale for long-range genomic prediction. bioRxiv, 2025. doi: 10.64898/2025.12.22.695963.

12. Hugo Dalla-Torre, Liam Gonzalez, Javier Mendoza-Revilla, Nicolas Lopez Carranza, Adam Henryk Grzywaczewski, Francesco Oteri, Christian Dallago, et al. Nucleotide Transformer: building and evaluating robust foundation models for human genomics. Nature Methods, 22(2):287–297, 2025. doi: 10.1038/s41592-024-02523-z.

13. Žiga Avsec, Natasha Latysheva, Jun Cheng, Guido Novati, et al. Advancing regulatory variant effect prediction with Al-phaGenome. Nature, 649:1206–1218, 2026. doi: 10.1038/s41586-025-10014-0.

14. Gonzalo Benegas, Carlos Albors, Alan J Aw, Chengzhong Ye, and Yun S Song. A DNA language model based on multispecies alignment predicts the effects of genome-wide variants. Nature Biotechnology, 43(12):1960–1965, 2025. doi: 10.1038/s41587-024-02511-w.

15. Max Schubach, Thorben Maass, Lusiné Nazaretyan Sebas-tian Röner, and Martin Kircher. CADD v1.7: using protein language models, regulatory CNNs and other nucleotide-level scores to improve genome-wide variant predictions. Nucleic Acids Research, 52(D1):D1143–D1154, 2024. doi: 10.1093/nar/gkad989.

16. Martin Kircher, Daniela M Witten, Preti Jain, Brian J O’Roak, Gregory M Cooper, and Jay Shendure. A general framework for estimating the relative pathogenicity of human genetic variants. Nature Genetics, 46(3):310–315, 2014. doi: 10.1038/ng.2892.

17. Vihaan Mathur and Ravi Sachidanandam. Benchmarking DNA foundation models: Biological blind spots in Evo2 variant-effect prediction. bioRxiv, 2026. doi: 10.64898/2026.03.10.710786.

18. Haonan Feng, Lang Wu, Bingxin Zhao, Chad Huff, Jian-jun Zhang, Jia Wu, Lifeng Lin, Peng Wei, and Chong Wu. Benchmarking DNA foundation models for genomic and genetic tasks. Nature Communications, 16:10780, 2025. doi: 10.1038/s41467-025-65823-8.

19. Thomas Dooms, Nicholas K Wang, and Michael Pearce. Covariance-based sequence pooling. https://www.goodfire.ai/research/covariance-pooling, 2026.

20. Michael T Pearce, Thomas Dooms, Alice Rigg, Jose Oramas, and Lee Sharkey. Bilinear MLPs enable weight-based mechanistic interpretability. In The Thirteenth International Conference on Learning Representations, 2025. URL https://openreview.net/forum?id=gI0kPklUKS.

21. Thomas Dooms and Ward Gauderis. Finding manifolds with bilinear autoencoders. In Mechanistic Interpretability Workshop at NeurIPS 2025, 2025. URL https://openreview.net/forum?id=ybJXIh4vcF.

22. Baiyu Lu, Xueshen Liu, Po-Yu Lin, and Nadav Brandes. Genomic heterogeneity inflates the performance of variant pathogenicity predictions. bioRxiv, 2025. doi: 10.1101/2025.09.05.674459.

23. Gregory M Findlay, Riza M Daza, Beth Martin, Melissa D Zhang, Anh P Leith, Molly Gasperini, Joseph D Janizek, Xingfan Huang, Lea M Starita, and Jay Shendure. Accurate classification of BRCA1 variants with saturation genome editing. Nature, 562:217–222, 2018. doi: 10.1038/s41586-018-0461-z.

24. Sounak Sahu et al. Saturation genome editing-based clinical classification of BRCA2 variants. Nature, 638:538–545, 2025. doi: 10.1038/s41586-024-08349-1.

25. Julianne S Funk et al. Deep CRISPR mutagenesis characterizes the functional diversity of TP53 mutations. Nature Genetics, 57(1):140–153, 2025. doi: 10.1038/s41588-024-02039-4.

26. Daniel R Tabet et al. The functional landscape of coding variation in the familial hypercholesterolemia gene LDLR. Science, 391(6787):eady7186, 2026. doi: 10.1126/science.ady7186.

27. Mitja I Kurki, Juha Karjalainen, Priit Palta, Timo P Sipilä, Kati Kristiansson, Kati M Donner, Mary P Reeve, Hannele Laivuori, Mervi Aavikko, Mari A Kaunisto, et al. Finngen pro-vides genetic insights from a well-phenotyped isolated population. Nature, 613(7944):508–518, 2023. doi: 10.1038/s41586-022-05473-8.

28. Sean V Tavtigian, Marc S Greenblatt, Steven M Harrison, Robert L Nussbaum, Snehit A Prabhu, Kenneth M Boucher, and Leslie G Biesecker. Modeling the ACMG/AMP variant classification guidelines as a Bayesian classification framework. Genetics in Medicine, 20(9):1054–1060, 2018. doi:10.1038/gim.2017.210.

29. Liv Gorton, Nicholas Wang, Nam Nguyen, Myra Deng, Eric Ho, Daniel Balsam, and Thomas McGrath. Interpreting Evo 2: Arc Institute’s next-generation genomic foundation model. https://www.goodfire.ai/research/interpreting-evo-2, 2025.

30. Onkar Gujral, Mihir Bafna, Eric Alm, and Bonnie Berger. Sparse autoencoders uncover biologically interpretable features in protein language model representations. Proceedings of the National Academy of Sciences, 122(34): e2506316122, 2025. doi: 10.1073/pnas.2506316122.

31. Vikas Pejaver, Alicia B Byrne, Bing-Jian Feng, Kymberleigh A Pagel, Sean D Mooney, Rachel Karchin, Anne O’Donnell-Luria, Steven M Harrison, Sean V Tavtigian, Marc S Greenblatt, Leslie G Biesecker, Predrag Radivojac, Steven E Brenner, and ClinGen Sequence Variant Interpretation Working Group. Calibration of computational tools for missense variant pathogenicity classification and ClinGen recommendations for PP3/BP4 criteria. The American Journal of Human Genetics, 109(12):2163–2177, 2022. doi: 10.1016/j.ajhg.2022.10.013.

32. Helen H Hobbs, David W Russell, Michael S Brown, and Joseph L Goldstein. The LDL receptor locus in familial hypercholesterolemia: mutational analysis of a membrane protein. Annual Review of Genetics, 24:133–170, 1990. doi: 10.1146/annurev.ge.24.120190.001025.

33. Howard J Worman. Nuclear lamins and laminopathies. The Journal of Pathology, 226(2):316–325, 2012. doi: 10.1002/path.2999.

34. Homa Tajsharghi and Anders Oldfors. Myosinopathies: pathology and mechanisms. Acta Neuropathologica, 125(1): 3–18, 2013. doi: 10.1007/s00401-012-1024-2.

35. Andreas C Joerger and Alan R Fersht. The p53 pathway: origins, inactivation in cancer, and emerging therapeutic approaches. Annual Review of Biochemistry, 85:375–404, 2016. doi: 10.1146/annurev-biochem-060815-014710.

36. Anthropic. Introducing Sonnet 4.6. https://www.anthropic.com/news/claude-sonnet-4-6, 2026.

37. Anthropic. Introducing Opus 4.6. https://www.anthropic.com/news/claude-opus-4-6, 2026.

38. ClinGen Sequence Variant Interpretation Working Group. ClinGen sequence variant interpretation recommendation for PM2 — version 1.0. https://clinicalgenome.org/site/ assets/files/5182/pm2_-_svi_recommendation_-_approved_sept2020.pdf, 2020.

